# Competition and coevolution drive the evolution and the diversification of CRISPR immunity

**DOI:** 10.1101/2021.11.12.468349

**Authors:** Martin Guillemet, Hélène Chabas, Antoine Nicot, François Gatchich, Enrique Ortega-Abboud, Cornelia Buus, Lotte Hindhede, Geneviève M. Rousseau, Thomas Bataillon, Sylvain Moineau, Sylvain Gandon

**Affiliations:** CEFE, CNRS, Univ Montpellier, EPHE, IRD, Montpellier, France; Institute of Integrative Biology, Department for Environmental System Science, ETH Zurich, 8092 Zurich, Switzerland; Bioinformatics Research Centre, Aarhus University, Aarhus C, Denmark; Département de biochimie, de microbiologie et de bio-informatique, Faculté des sciences et de génie, Université Laval, Québec City, G1V 0A6, Canada; Groupe de recherche en écologie buccale, Faculté de médecine dentaire, Université Laval, Québec City, G1V 0A6, Canada; Félix d’Hérelle Reference Center for Bacterial Viruses, Université Laval, Québec City, G1V 0A6, Canada

## Abstract

The diversity of resistance challenges the ability of pathogens to spread and to exploit host populations [1–3]. Yet, how this host diversity evolves over time remains unclear because it depends on the interplay between intraspecific competition among host genotypes and coevolution with pathogens. Here we study experimentally the effect of coevolving phage populations on the diversification of bacterial CRISPR immunity across space and time. We demonstrate that the negative-frequency-dependent selection generated by coevolution is a powerful force that maintains host resistance diversity and selects for new resistance mutations in the host. We also find that host evolution is driven by asymmetries in competitive abilities among different host genotypes. Even if the fittest host genotypes are targeted preferentially by the evolving phages they often escape extinctions through the acquisition of new CRISPR immunity. Together, these fluctuating selective pressures maintain diversity, but not by preserving the pre-existing host composition. Instead, we repeatedly observe the introduction of new resistance genotypes stemming from the fittest hosts in each population. These results highlight the importance of competition on the transient dynamics of host-pathogen coevolution.

## 1 Introduction

Coevolution is thought to be a powerful evolutionary force at the origin of biological diversity [4–6]. Negative-frequency-dependent selection generated by coevolution can promote the emergence and the maintenance of genetic diversity in interacting species [5, 7, 8]. On the other hand, maintenance of genotype diversity is also affected by intrinsic differences in competitive abilities among genotypes. If this asymmetric competition is strong it can lead to the exclusion of less competitive genotypes and to a drop in diversity. The interplay between coevolution and competition has been explored theoretically with models based on the “kill-the-winner” hypothesis which explicitly accounts for the influence of phages on diverse host communities [9–11]. This framework, however, is meant to describe the ecological dynamics of interacting host species. Several experimental studies have explored the influence of phages on interspecific competition among different bacterial species [12–15] but intraspecific competition is harder to monitor. Studying the interplay between competition among host genotypes and coevolution with pathogens is particularly challenging within a host species because it requires detailed knowledge of the genetic determinants of the specificity of the host-virus interaction to track the dynamics of different host genotypes [16].

Here, we track the coevolutionary dynamics of CRISPR immunity of the bacterial species *Streptococcus thermophilus* with its lytic phage 2972. This model system offers unique opportunities to explore the microevolutionary processes driven by competition among different bacteria and antagonistic coevolution between bacteria and their viral pathogens. In *S. thermophilus*, coevolution with phage 2972 is mainly driven by two (type II-A) CRISPR-Cas loci (CR1 and CR3) which allow the bacteria to incorporate 30-bp DNA sequences (spacers) from the genome of an infecting phage in the CRISPR array [17–20]. After transcription, each spacer RNA is used as a guide by Cas9 to target and cleave the corresponding target sequence (the protospacer) in the phage genome, thereby halting virus replication and reducing its titer. Phages can escape CRISPR immunity via mutations in the protospacers which avert recognition by the Cas complex. These mutations have been shown to be particularly effective at escaping immunity when they are located at specific positions in the proto-spacers like the PAM (protospacer-adjacent motif) or the seed [21, 22]. Crucially, the sequencing of the CRISPR array of the populations of bacteria and the whole-genome sequencing of the populations of phages allowed us to fully characterize the specificity of the infection network, without any phenotypic assays. Here we focus on the CRISPR array of the CR1 locus which has been shown to be the most active of the CRISPR loci of S. *thermophilus* against phage 2972 [18].

To study how host diversity affects the dynamics of CRISPR immunity, we designed a short-term coevolution experiment (pictured in Figure 1) where we followed the evolution of CRISPR immunity in the absence of phages (treatment A), in the presence of an initially monomorphic population of phages (treatment B), in the presence of an initially polymorphic population of phages (treatment C). We started each culture with a mix of 17 different bacterial strains in equal frequencies: one strain was fully sensitive to the wild-type lytic phage 2972 (strain DGCC 7710) and each of the remaining sixteen strains carried a distinct single-spacer resistance in the CRISPR array at the CR1 locus (Table S1). These strains were obtained from a previous study after exposing the susceptible strain DGCC 7710 to the phage 2972, leading to the spontaneous acquisition of a single additional spacer targeting distinct protospacers in the phage genome [23]. Crucially, the bacteria may have acquired additional mutations in the bacterial genome outside of the CRISPR loci during this selection procedure. We carried out whole genome sequencing of all the strains to identify these mutations (Table S3 and in the Supplementary Data). A previous study demonstrated that the ability of phage 2972 to escape CRISPR immunity differs among the sixteen resistant strains [23].

**Figure 1:**
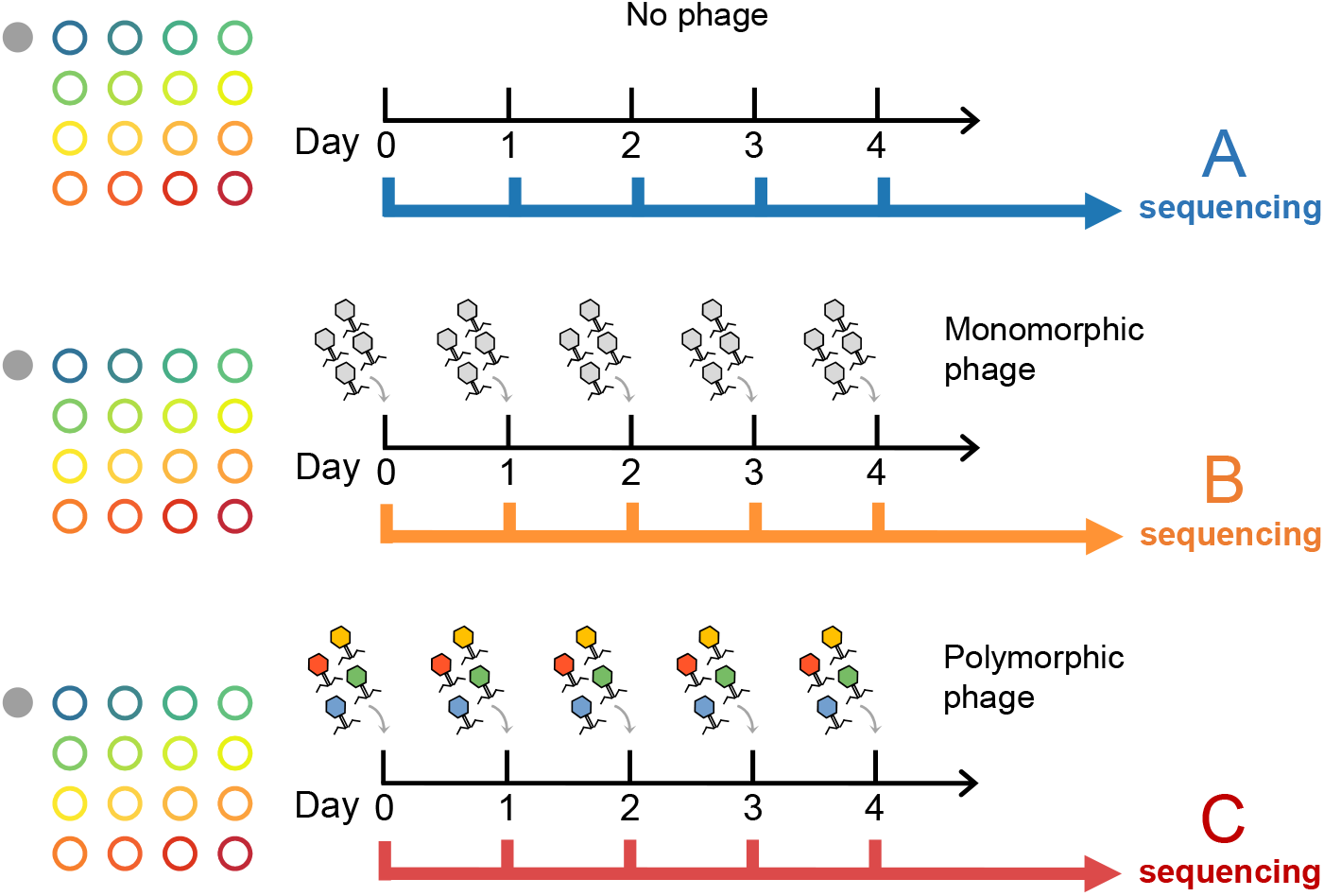
The three treatments of our coevolutionary experiment. Bacterial cultures were inoculated with a mix of 17 different strains in equal frequencies: one strain (filled grey circle) was susceptible to the wild-type lytic phage 2972 and the remaining sixteen strains (empty colored circles) carrying a distinct single-spacer resistance in the CRISPR 1 (CR1) locus. (A) daily transfers of 1% of the bacterial culture with no exposure to phages, (B) daily transfers of 1% of the bacterial culture with inoculation of 10^5^ phages at each transfer sampled from a monomorphic population of the wild-type phage, (C) daily transfers of 1% of the bacterial culture with inoculation of 10^5^ phages at each transfer sampled from a polymorphic phage population. This polymorphic phage population is a mix of sixteen escape variants that were previously selected to escape each of the sixteen CR1 resistance of the polymorphic population of bacteria.

For each of the three experimental treatments, we transferred 1% of each replicate culture to a fresh medium for 4 consecutive days. In the absence of phage (treatment A), the change in the relative frequency of the different host genotypes informed us about the competitive abilities of the 17 bacterial strains. This treatment allowed us to evaluate the ability to maintain diversity on the CRISPR locus in the absence of selection for resistance (Figure S1). If some strains are more competitive they are expected to outgrow the others and induce a rapid drop of diversity. The two other treatments allowed us to follow the interplay between competition and antagonistic interactions with phages on the evolution of the bacteria. At the beginning of each transfer, we added 10^5^ phages from a monomorphic or a polymorphic phage populations (Treatments B and C, respectively). The monomorphic phage population was obtained from the amplification of the wild-type phage 2972 which infects only the sensitive host strain (about 6% of the host population at the onset of our experiment). In the polymorphic phage population, we used a mix of sixteen escape phage variants (phage cocktail) that were previously selected to escape each of the sixteen CRISPR CR1 resistances of the polymorphic population of bacteria [23] (Table S2). This recurrent inoculation of phages at each transfer was used to maintain a minimal amount of phage in treatments B and C. As pointed out below, this immigration of phages did not prevent phage adaptation and coevolution with the host.

To monitor the demography and evolution of bacteria we used spacers as barcodes and sequenced the 5’-end of the CRISPR array of the CR1 locus of the bacteria (see Methods). This sequencing strategy allowed us to identify the emergence and the spread of additional resistant strains with new spacers in the CRISPR array [24]. To monitor the evolution of the phage populations we used whole genome sequencing in the treatments exposed to the virus (Treatments B and C) to identify new mutations and estimate their frequencies.

## 2 Results and Discussion

### 2.1 Phage diversity drives infection dynamics

The treatments had major effects on both the bacteria and the phage densities (Figure 2). The monomorphic phage treatment had a limited impact on bacterial growth the first day but led to a massive phage epidemic on the second day, marked by a drop in host density and an increase in the viral pathogen density. In contrast, the polymorphic phage treatment immediately led to a viral outbreak on the first day. Yet, under all phage treatments the bacterial populations eventually recovered and by day 4 they reached a density close to the no-phage treatment (Figure 2a).

**Figure 2:**
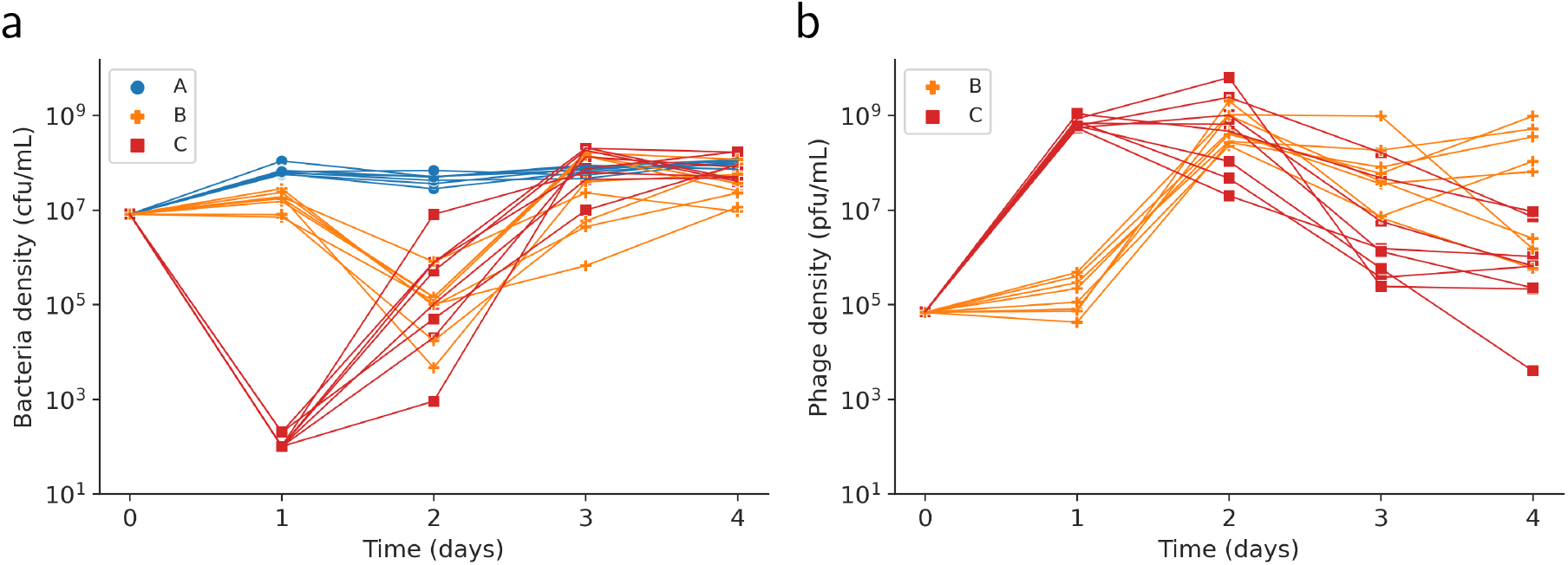
Demography of bacteria and phages: (a) density of bacteria and (b) density of phages in the three experimental treatments. All replicates are shown, 7 for the control (treatment A) and 8 for the two phage treatments (treatments B and C). Blue points show the data in the absence of phages, while orange and red show the data for the monomorphic and polymorphic phage treatments, respectively.

### 2.2 Evolution and diversification of CRISPR immunity

To monitor the evolutionary dynamics of bacteria, we tracked the diversity of CRISPR immunity at the CR1 locus and estimated the frequency *h_i_* of each resistance genotype *i* in the population. We computed the effective number of host genotypes [25] across time for each replicate (Figure 3). Strikingly, the effective number of host genotypes dropped very fast in the treatment without phage and remained very low until the end of the experiment. Phage treatments had a significant effect on the mean effective number of host genotypes (see Methods)(Day 1: F_2,20_ = 1431, *P* = 2.6*e* – 22; Day 4: F_2,20_ = 7.80, *P* = 3.1*e* – 3). Compared to the treatment without phage at Day 1 (effective number of genotype and 95% CI: 8.90 [8.72, 9.10]), exposure to a monomorphic phage population initially led to a faster drop in diversity (6.91 [6.74, 7.08]), but exposure to a polymorphic phage treatment maintained a high level of diversity (14.88 [14.63, 15.12]). Both phage treatments led to the maintenance of more diversity at the end of the experiment than the treatment without phage (Day 4, no phage: 1.96 [1.92, 1.99], monomorphic phage: 3.66 [2.79, 4.52]; polymorphic phage: 5.33 [3.78, 6.98]). The maintenance of diversity in host populations exposed to phages supports the idea that coevolution can drive the diversification of host populations [2, 4, 10, 26]. The variation in the dynamics of diversity among replicate populations exposed to phages illustrates the impact of demographic stochasticity on this coevolutionary dynamics, particularly after demographic bottlenecks caused by viral epidemics.

**Figure 3:**
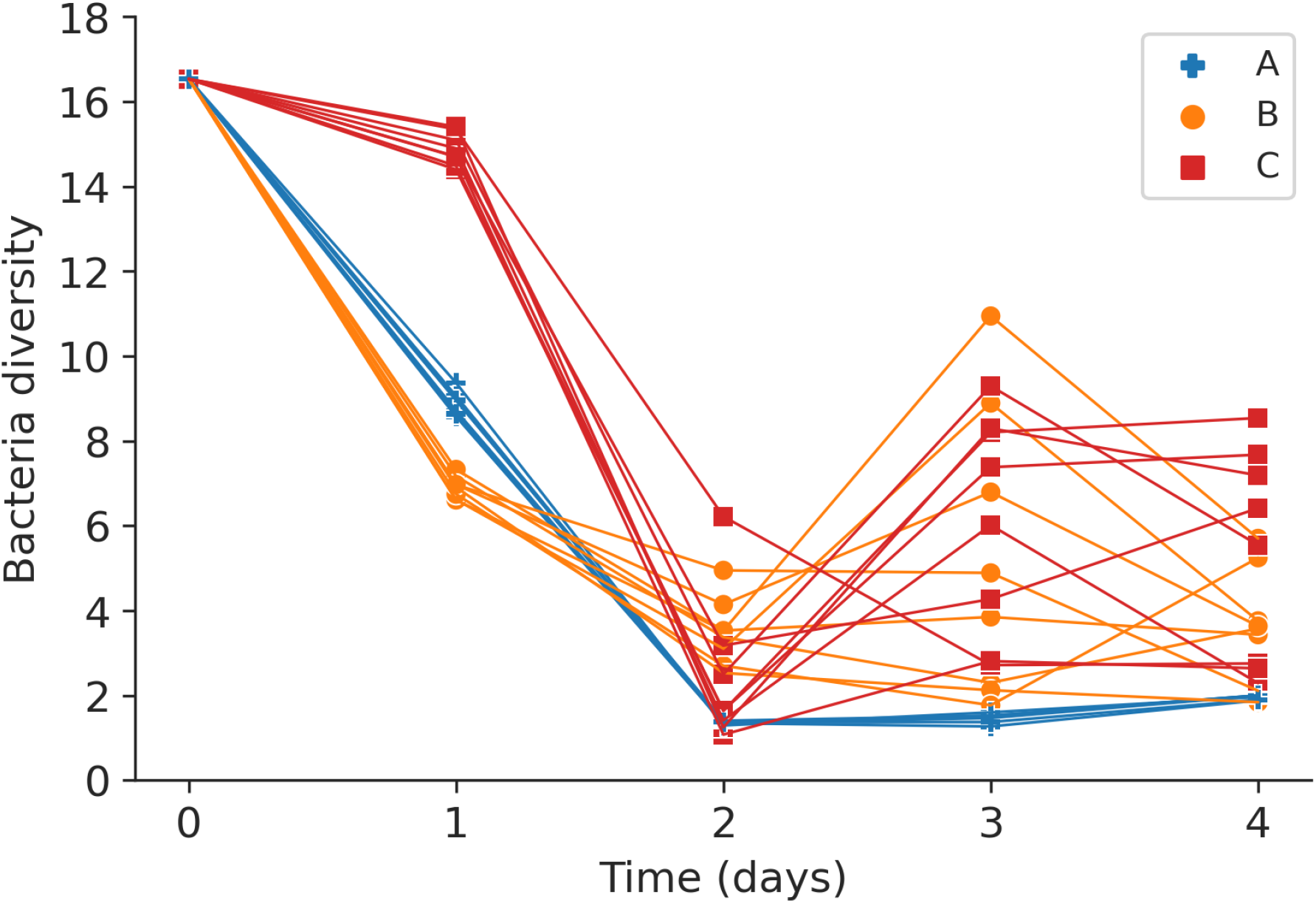
Dynamics of the diversity of CRISPR immunity. Dynamics of CRISPR locus diversity computed with the effective number of host genotypes: 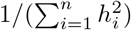. Blue points show the data in the absence of phages, orange and red show the data for the monomorphic and the polymorphic phage treatments, respectively.

Next, to better understand what drives the dynamics of CRISPR diversity we examined the competition between the different bacterial strains using modified Muller plots that provide a description of both the changes in density and in the genetic composition for each replicate population of bacteria (Figure 4). All the replicates followed very similar dynamics in the treatment without phage (Figure 4): one of the strain (indicated in red, strain 31725) outcompeted the other strains and nearly reached fixation by day 2, but another strain (indicated in green, strain 16236) increased in frequency towards the end of the experiment. These results indicate major differences in competitive abilities among strains. Interestingly, the fitter strain (in red) is not the phage-sensitive wild-type strain but one of the sixteen resistant strains (Figure S1). Whole genome sequencing of the seventeen strains used at the beginning of the experiment revealed the existence of other mutations across the bacterial genome outside the CRISPR locus (Table S3). These mutations were acquired during the selection process that led to the natural acquisition of a new spacer on the CR1 locus [27]. For instance, the “red” strain has eight unique non-synonymous mutations in different genes. In contrast, the sequencing of the “green” strain revealed only two unique synonymous mutations in different genes. The competitive ability of the “green” strain is also more puzzling because this strain was initially less fit and only increased in frequency towards the end of the experiment. A more detailed analysis of the contribution of each of these mutations on the competitive ability of the strains falls beyond the scope of this study. But these highly consistent measures of fitness among replicates in the treatment without phage allowed us to study how competition affects the coevolutionary dynamics in the populations exposed to phages.

**Figure 4:**
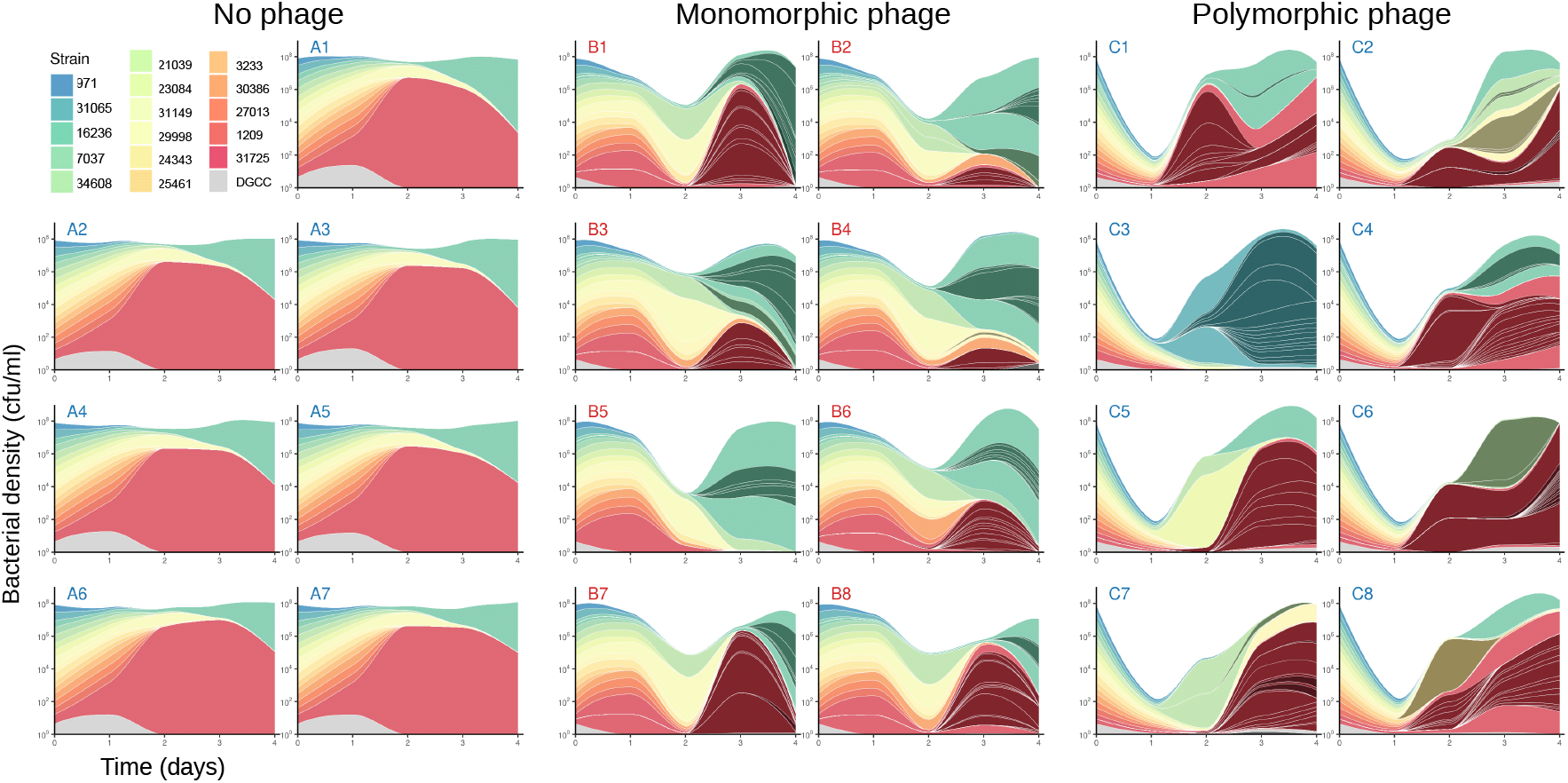
Host populations resist phages through the diversification of the CR1 locus. Modified Muller plots show the dynamics of different host genotypes in each replicate of the three experimental treatments as indicated above each graph (‘A’ for the no phage control, ‘B’ for the monomorphic phage treatment and ‘C’ for the polymorphic phage treatment). The total height for each day shows the bacterial density (in cfu/ml) on a log scale, and the different colors show the proportion of the different host genotypes at each time point on a linear scale. The 17 strains that were added on day 0 (including the Wild-Type in grey) are shown in the legend. The blue-to-red color scale ranks the strains according to their initial fitness as detailed in figure S1. A darker colored strain seen in later days stemming from inside one of the 17 original strains shows the acquisition of a new spacer. An even darker color represents strains with 2 additional spacers (2 new spacers is the maximum represented here). When there are several parallel acquisitions of new spacers, the new strains are separated with white lines. The lines are smoothed between each day.

Figure 4 shows how phages affect both the density of bacteria and the evolution of CRISPR immunity. As expected from Figure 3, the presence of phages maintains a higher number of strains. More specifically, we observe the emergence of several new resistant strains that carry up to three additional spacers in the CRISPR array, which are indicated by dark colors in Figure 4. Interestingly, in all treatments exposed to phages, almost all the bacterial populations end up being dominated by lineages that are descendants of the two most competitive strains identified in the absence of phages (Figure S2). In other words, the increase in diversity observed at the end of the experiment in the populations exposed to phages (Figure 3) is not due to the initial diversity being restored, but to new resistance genotypes that arose via the acquisition of new spacers in the CRISPR array of the winners of the competition among bacterial strains (Figure S3). It is unlikely that the per capita rate of acquisition of new spacers differs among the different bacterial strains. Indeed, none of the mutations found in these strains were in the genes known to control the adaptation step of type II-A CRISPR-Cas system (i.e. *cas1, cas2, csn2, cas9*). Variation in the densities of bacteria provides a more parsimonious explanation for the faster acquisition of new spacers in the winners of the competition. Since the winners of the competition were more abundant, they were also more likely to acquire new spacers.

Importantly, the comparison among replicate populations revealed very different dynamics in the presence of phages. To study this variation, we measured the amount of genetic differentiation among replicate populations within each treatment (Figure S4). Complementary measures of host differentiation (*F_ST_* and *D*) allowed us to quantify the changes in population composition due to drift and selection among replicates (see Methods). As expected, differentiation remained very low in the treatment without phages because all replicates displayed very similar dynamics. In contrast, exposure to phages led to the acquisition of distinct spacers in different replicates, which led to a rapid increase in differentiation among host populations. This is particularly noticeable right after the massive demographic bottleneck that took place after the first day in the treatment exposed to a polymorphic phage population.

Another way to demonstrate the influence of phages on bacterial evolution is to detect the presence of negative-frequency-dependent selection. As expected, in the absence of phages the change in strain frequency between time *t* and *t* + 1 is independent of strain frequency at time *t* (Figure 5a). Exposure to phages, however, yields a strongly negative relationship between these two quantities (the presence of phages has a highly significant effect on the slope of the regression line in both the monomorphic and the polymorphic phage treatments, see Methods), which indicates that more frequent strains tend to be selected against because they are preferentially targeted by phages (Figures 5b and 5c). All these results confirm the expected impact of viral pathogens on the diversification of host resistance [26, 28] and highlight the relevance of the “kill-the-winner” hypothesis [9, 10].

**Figure 5:**
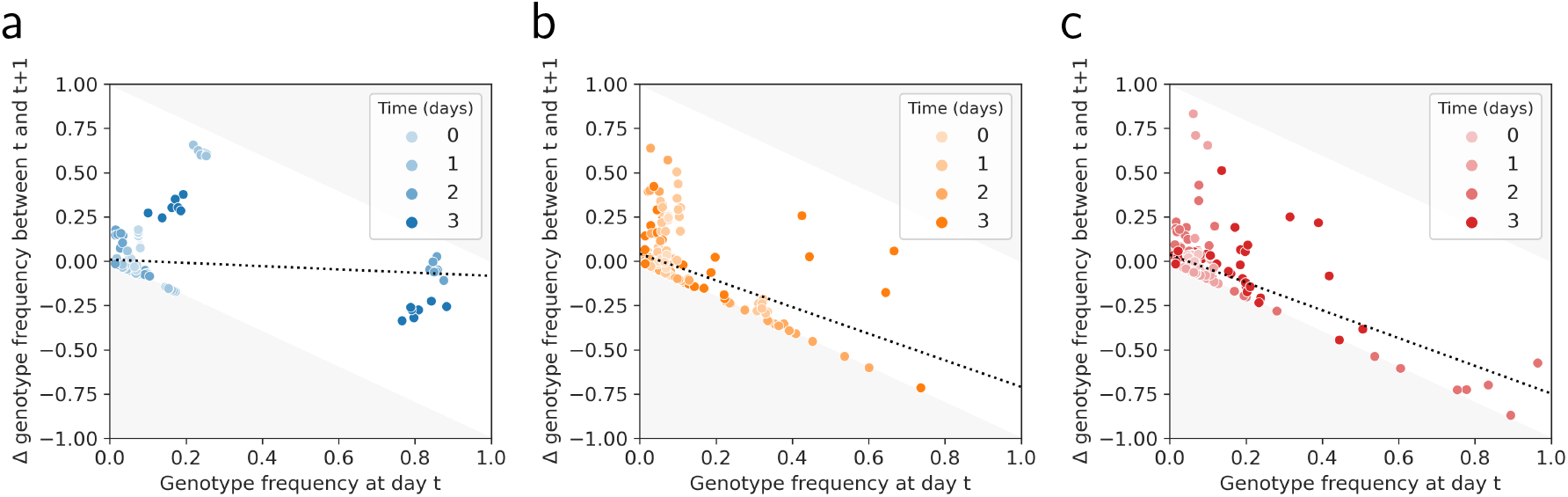
Phages induce negative frequency dependent selection. Evidence for Negative Frequency Dependent Selection (NFDS) in the two phage treatments (b,c) compared to the control (a). In every replicate, the change in frequency Δ of a genotype between day *t* and *t* + 1 is plotted according to its frequency at day *t*. A linear regression is plotted in each panel to highlight the NFDS (or lack thereof in the control). A significantly negative slope indicates that more frequent genotypes are counter-selected and tend to decrease in frequency the following day (see Methods). The light grey area refers to unfeasible changes in frequency. Blue points show the data in the absence of phages, orange and red dots show the data for the monomorphic and polymorphic phage treatments, respectively.

### 2.3 Phage coevolution across space and time

The sequencing of the phage populations revealed the emergence and the spread of many mutations across the phage genome (Figure S5). Most of these mutations were located in the protospacer regions targeted by CRISPR immunity and particularly in the PAM or the seed of protospacers (Figure S6). These mutations are expected to be strongly beneficial as they allow the virus to escape CRISPR immunity [21]. Knowing the genetic specificity of CRISPR immunity allows us to assign phenotypic effects to these mutations without any additional experimental measures. We combined sequencing data from the bacteria and the phages to compute the mean fitness 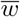 of each phage population using:

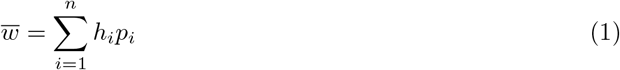

where *n* is the total number of host genotypes, *h_i_* is the frequency of host genotype *i* and *p_i_* is the frequency of phage variants that can infect genotype *i*. Here, the mean fitness measures the mean fraction of the host population available to a randomly sampled virus in the phage population. This *in silico* measure of phage mean fitness provides a powerful way to estimate phage adaptation to contemporaneous host populations (when phage and bacteria frequencies are sampled in the same replicate and at the same point in time) but also across space and time [6, 26].

Measures of phage adaptations across all time points revealed a striking pattern where levels of phage adaptation are maximal against host populations from the recent past (Figure 6). In contrast, the degree of phage adaptation drops very rapidly against bacteria from the future in both phage treatments. This pattern is precisely the one expected under the rapid coevolution dynamics that are predicted to emerge in coevolutionary models [6, 29–31]. The particularly rapid drop of phage mean fitness when matched against bacteria from the future shows how quickly bacteria are able to develop new resistance to the phages. This is consistent with the intrinsic asymmetry inherent to CRISPR specificity: bacteria have access to hundreds of different protospacers from the phage genome [21] allowing them to raise a diverse and distributed immune defense to the phage population at once [32]. In contrast, only mutations in the targeted viral genomic region (Figure S6) can provide an effective way for phages to escape CRISPR immunity, and only against one resistance (one spacer) at a time.

**Figure 6:**
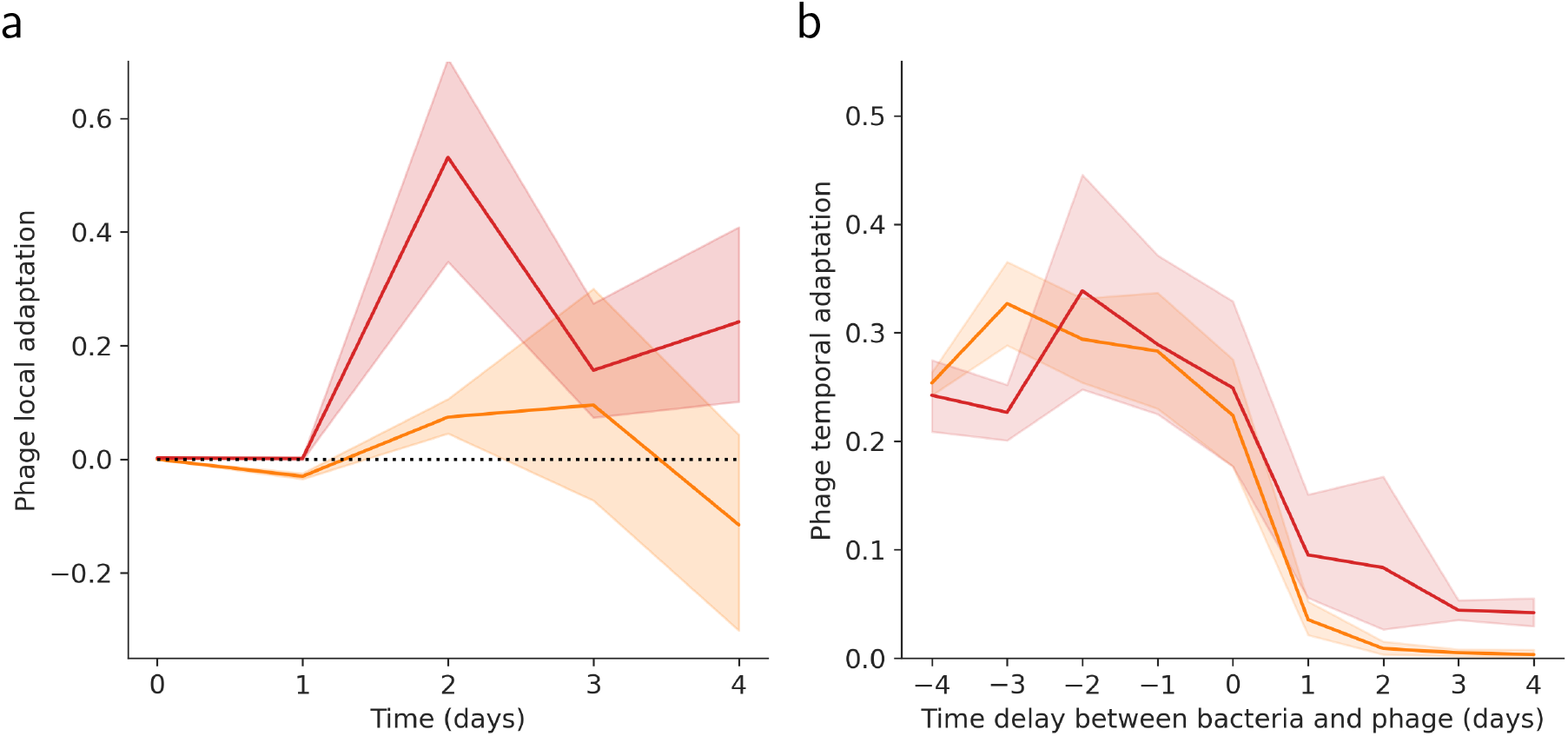
Phage adaptation across space and time. Evidence for (a) local and (b) temporal adaptation to CRISPR immunity in the two phage treatments. Local adaptation measures the amount of phage adaptation to sympatric host populations (same replicate) relative to allopatric host populations (different replicates) for each treatment at different points in time (see Methods). Temporal adaptation measures, for a given replicate, the mean fitness of the phage population to host sampled at different points in time. A positive (negative) time delay between bacteria and phage indicates that the bacteria is sampled in the future (past) relative to the sampling point of the phage (see Methods). Orange and red show the average level of phage adaptation for the monomorphic and polymorphic phage treatments, respectively. The shaded areas show the 95% confidence interval computed using the Jackknife method (see Methods). The horizontal dotted line in (a) shows the position of the zero, where there is no local adaptation.

Measures of mean fitness across space allowed us to compute phage local adaptation to determine if the phage is more adapted to sympatric (same replicate) than to allopatric (different replicate) host populations (see Methods) [29, 33]. Figure 6 shows the buildup of local adaptation across time in the two phage treatments. Interestingly, local adaptation remains very low in the treatment with the monomorphic phage population. In contrast, we detect a strong pattern of phage local adaptation in the treatment with a polymorphic phage population. In particular, phage local adaptation is extremely strong (0.53[±0.18]) at day 2 which coincides with the time at which host differentiation is maximal. Indeed, phage local adaptation can only occur when the composition of sympatric host populations differ substantially from allopatric host populations.

The dynamics of differentiation varied between the two phage treatments. Even if we detect significant differentiation among replicate populations in the treatment with the monomorphic phage population, the *Q_ST_* which measures phenotypic differentiation (see Methods) remains very low (Figure S7) because most escape mutations occur on the same protospacer (Figure S5). Indeed, the high number of escape mutations we observe (in the monomorphic phage treatment) in the terminal region of the phage genome (around 31kb, Figure S5) can be explained by the presence of the protospacer associated with the “red” resistant host (strain 31725). Selection to escape this resistance was very intense in this treatment (because this resistance strain was the most frequent) and each of these mutations correspond to different solutions leading to the same escape phenotype. In contrast, in the treatment with a polymorphic phage population, both *F_ST_* and *Q_ST_* are increasing on day 2 after the divergence of bacterial populations (Figure S7) and the distribution of escape mutations is more evenly distributed across the phage genome (Figure S5). Note, however, that the speed of phage adaptation seems too low to catch up with the build up of CRISPR immunity. Initial diversity in the polymorphic treatment yields faster adaptation but the number of phage mutations in protospacers stops increasing by day 1 (Figure S8). In the monomorphic treatment, the saturation in the number of mutations in protospacers is delayed (Figure S8). The drop in local adaptation with time is consistent with the overall drop in phage density we observed in most phage populations (Figure 2b). This suggests that the phages are losing the coevolutionary arms race with their hosts which is in line with previous studies showing that CRISPR immunity often yields phage extinction in this system [34–36]. Besides, we found some evidence that evolution of new resistance may also be due to the second active CRISPR-Cas system (CR3) in this host in which we detected spacer acquisition starting at day 3 or 4 (Table S4). Accounting for evolution at the CR3 locus when estimating phage fitness magnifies the drop of mean fitness of phage populations (Figure S9). The recurrent introduction of ancestral phages at the beginning of each transfer was used to avoid phage extinctions in treatments B and C. This recurrent immigration probably had a limited impact on the coevolutionary dynamics in our short-term experiment because immigrants were unable to infect bacteria that rapidly acquired new resistances. Yet, migration could have more impact on long-term persistence as it would allow phages to recolonise susceptible host populations.

### 2.4 Host competition governs the coevolution-driven diversification

We can track the dynamics of phage adaptation across space and time but can we predict the speed at which the phage escapes the phage-resistant strains? The speed of adaptation is governed (i) by the rate of mutation, which has been shown to vary among protospacers in a previous experiment [23], (ii) by the strength of selection associated with the ability to escape CRISPR immunity against a specific protospacer and (iii) by the fitness cost of these escape mutations. Because the fitness cost of these mutations has been shown to be a poor predictor of the durability of CRISPR-Cas immunity [23] we focus on the first two points. In the treatment with a polymorphic phage population, the rate of mutation is not limiting because the mutations against the 16 original spacers are pre-existing. In this phage treatment, as expected, we do not find a correlation between the speed of phage adaptation and the rate of escape mutation for different protospacers (Pearson’s *r* = −0.26, *P* = 0.41) (Figure S10). In contrast, the speed of adaptation is governed by the competitive ability of the different resistant strains (Pearson’s *r* = 0.83, *P* = 7.5e-5). Indeed, this competitive ability is a good predictor of the abundance of each resistant strain and, consequently, a good predictor of the fitness benefit associated with the ability to exploit these resistant strains. Interestingly, we obtain a very similar pattern in the monomorphic phage treatment (no correlation with the mutation rate: Pearson’s *r* = −0.02, *P* = 0.41; strong correlation with competitive ability: Pearson’s *r* = 0.95, *P* = 3.4e-8). These results indicate that phage mutation is not limiting and phage adaptation is mostly driven by the more abundant (i.e. the more competitive) phage-resistant strains of bacteria.

## 3 Conclusion

Our short-term evolution experiment demonstrates that the coevolutionary battle taking place between bacteria and phages is a potent evolutionary force driving the rapid diversification of interacting populations. The presence of phages generates strong negative-frequency-dependent selection, which prevents the loss of diversity of CRISPR immunity. This is consistent with the “kill-the-winner” hypothesis [9, 10] which states that viruses can maintain the host diversity. Similar conclusions were reached from studies that explored the interplay between interspecific competition and coevolution with phages [13, 15]. But here, we could track the emergence of new resistance mutations (new spacers in the CRISPR array) and these new mutations are not equally distributed among the bacterial strains present initially. Indeed, we see that the initial host diversity vanished rapidly (Figures S3) and in all but one replicate population exposed to phages, the bacterial population at day 4 is dominated by strains that descend from the most competitive strains (the “winners”) identified in the control (Figures 3 and S2). To understand these results it is important to recall that host adaptation results from both the selection imposed by phages at the CRISPR locus and the selection imposed on the rest of the bacterial genome. The recurrent bottlenecks in the host population size induced by phage infections may lead to a faster fixation of new mutations. Even if these additional mutations are expected to be often deleterious [37], their effects on fitness will vary and introduce variation in competitive abilities among strains [37, 38]. In the absence of phages, fitter host genotypes consistently outcompete other strains. In the presence of the phage, viral adaptation targets preferentially more abundant and competitive strains. But the evolution of CRISPR immunity allows the winners of the intraspecific competition to strike back after phage adaptation. Ultimately, this explains why diversity is generally maintained and originates from the descendants of the winners in populations exposed to phages. This feedback of competition on coevolutionary dynamics can also be discussed in the light of the recent “royal family model” [39]. In a classic version of the “kill-the-winner” framework, the most frequent host strain is preferentially targeted by the evolving population of pathogens and is driven to low frequency. Next, another host strain rises to high frequency and the cycle repeats. In the “royal family model” intrinsic asymmetries in competitive abilities imply that the newly rising host genotypes are likely to descend from the previously dominating genotypes. Our experimental results squarely fit within this framework as we can readily identify a royal family in the bacteria population which often derives from the more competitive strains (the red and green strains in Figure 4). Note, however, that our experiment also features the rise of a new royal family (strain 31065) in one population after a particularly strong demographic bottleneck (replicate C3 in Figures 4 and S2). Hence, the stochastic acquisition of new resistance may open up new opportunities for previously dominated strains of bacteria. As expected from the “royal family model” this evolutionary dynamics within the population of bacteria implies that there is also a royal family of phages, which is particularly adapted to the royal family of bacteria (Figure S11). We stress that our short-term experiment focuses on a very specific scenarios where (i) the initial diversity in the host population was manipulated artificially with equal frequency among different strains and no multiresistance to the phage and (ii) the initial diversity of the phage population was also manipulated experimentally (treatment B versus C). Yet, the distributions of CRISPR immunity and phage diversity are expected to buildup naturally after a phage epidemic and the network structure of strain diversity may be very different from the one used in our experiment [40]. Our work should be viewed as a first attempt to monitor coevolutionary dynamics experimentally and the relevance of the “royal family model” remains to be investigated in a more natural setting.

This short-term experiment demonstrates that ecological and evolutionary processes can take place on a similar time scale. A better understanding of the coevolution between CRISPR immunity and phages requires a more comprehensive theoretical framework considering the mutations involved in the interaction as well as in the rest of the genome. Current models of host-parasite coevolution neglect possible asymmetries in competitive abilities among host genotypes carrying the same number of resistance genes. However, our experiment shows that the accumulation of mutations in loci not involved in interactions with the phages can lead to a drop in the immune diversity after a local extinction of the phage population. This drop in resistance diversity is likely to facilitate the evolutionary emergence of the phages when new viruses are introduced in the population [2, 3]. We expect that this process may alter dramatically the coevolutionary dynamics studied with numerical simulations by [40]. The collapse of the diversity of CRISPR immunity in the absence of phage (or when phage are very rare) would shorten the duration of the Host Control Regime (HCR in periods in [40]) and would speed up the coevolutionary dynamics between phages and CRISPR immunity. At the larger spatial scale this succession of local phage extinction and rapid recolonisation could ensure the long-term coexistence of bacteria and phages in spatially structured environments.

## 4 Materials and Methods

### 4.1 Bacteria and bacteriophage strains

*S. thermophilus* DGCC 7710 and phage 2972 [41] were obtained from the Felix d’Hérelle Reference Center for Bacterial Viruses (www.phage.ulaval.ca). Sixteen derivative phage-resistant strains, each with an unique CRISPR spacer were generated previously [23] and sequenced to look for mutations outside of the CRISPR loci (Table S1). Similarly, sixteen phages carrying mutation to escape the resistance of these individual spacers were isolated after selection on each resistant bacteria (see list of protospacer sequences in Table S2) [23].

### 4.2 Experimental procedure

Prior to the experiment, the 17 bacterial strains were mixed and grown during 6 hours in LM17+CaCl_2_ (37 g/l of M17 (Oxoid) supplemented with 5 g/l of lactose and 10 mM of CaCl_2_). Then, the bacterial mix was transferred 1:100 into 10 ml of fresh LM17+CaCl_2_ (no phage treatment, 7 replicates), infected with 10^5^ wild-type 2972 phages (monomorphic phage treatment, 8 replicates) or infected with the mix of 10^5^ phages (polymorphic phage treatment, 8 replicates), then incubated at 42°C. We used only 7 replicates in the control because we were limited by the total number of replicates we could sequence using the Nextera XT 96 samples prep kit (see below). Every day (after 18 hours of incubation), 1% of the cultures were transferred into 10 ml of LM17+CaCl_2_, and 10^5^ phages were inoculated from the same population of phage (monomorphic or polymorphic) used at the beginning of the experiment. Following each transfer, the bacteria and phages from each replicate were separated by filtration (0.2 μm) and titrated as described in [23]. To guarantee that there was enough DNA for sequencing, the phages were reamplified once on susceptible host bacteria (i.e. DGCC 7710) over 5h (after full lysis of the bacteria), then DNA was extracted using the ZYMO Quick-DNA Miniprep plus kit. Note that this amplification step may have introduced some bias in phage mutation frequencies if some phage genotypes were more fit than others in an environment with only susceptible hosts.

### 4.3 Bacteria sequencing

The CRISPR-Cas CR1 locus was amplified through PCR (primers 5’-3’: AGTAAGGATTGACAAG-GACAGT; CCAATAGCTCCTCGTCATT) from the different populations from the three different treatments, the different replicates and the different time points. These PCR products were tagged using Nextera XT 96 samples prep kit and pooled before sequencing with Illumina MiSeq. The spacers were extracted from the sequences by searching for the flanking repeats allowing for a maximum of one mismatch. The spacers were then matched with their protospacers on the phage genome using Blast version 2.8.1 [42] and the protospacer database presented in the next section. After these steps, an average sequencing depth of around 95700 was obtained. A minimum identical word size of 10, and a 70% identity threshold was used. The top result of the search, if any, was used to replace the name of the spacer by the middle position of the protospacer in the phage genome. A frequency cutoff of 1% was used to optimize the quality of our dataset. The resulting frequencies of genotypes over time in each replicate are available in the Supplementary Data. We found that in the treatment with the monomorphic phage population there has been significantly more acquisition of spacers that were already present in the original 16 bacterial strains than the other 677 potential spacers (Chi-square test: chi2 obs=12.17, df =1, P =4.8e-4). This means that the spacers already present in the mix were acquired preferentially which may be due to DNA transfer among bacteria. The CRISPR-Cas CR3 locus was amplified through PCR (primers 5’-3’: GGTGACAGTCACATCTTGTCTAAAACG; GCTGGATATTCGTATAACATGTC) and migrated on 1.5% agarose gel to check for spacer acquisition. The samples with additional bands indicating the acquisition of an additional spacer are given in Table S4.

### 4.4 Phage sequencing

The phage DNA samples were sequenced (Illumina MiSeq) with 150-bp paired-end reads. Trimmomatic [43] was used to clean and trim the sequencing reads yielding an average sequencing depth of around 650, before mapping them on the reference genome using Bowtie2 [44]. The software FreeBayes [45] was then used to detect single-nucleotide polymorphism and the phage reference genome [41] was updated to include the SNPs with frequency > 0.45 in the initial mix, to distinguish these pre-existing mutations from the ones that arose during the experiment. The read mapping and the SNP detection were done a second time using this updated genome as reference. The resulting frequencies of phage mutations over time in each replicates are available in the Supplementary Data. To detect the protospacers in the phage genome, we looked for the CR1 specific PAM sequence ‘GGAA’ or ‘AGAA’ in both strands of this reference genome and found 693 occurrences (281 and 412 respectively for the two PAMs).

### 4.5 Fitness and adaptation estimates

We computed the mean phage fitness in a certain host population with equation (1). The frequencies of matching spacers and protospacer mutations are provided in the Supplementary Data. Our short-read sequencing data for the phages does not give linkage information between mutations so we need a linkage hypothesis to compute *p_i_* from the frequencies of escape mutation frequencies derived from whole-genome sequencing of phage populations. When the host resistance genotype *i* carried more than a single spacer we assumed that the genotype frequency of the phage variant able to infect host resistance genotype *i* was the product of the frequencies of the mutations on all the protospacers targeted by this set of spacers (i.e., we assumed linkage equilibrium among these mutations). To check the robustness of our results we computed phage fitness under the alternative assumption that escape mutations are fully linked (by setting to the frequency of phage mutations providing escape to the last spacer in the CRISPR locus). We observed a maximum of 2.7% difference between the measures of mean fitness of the phage in sympatric (same replicate) and contemporaneous (same time point) host populations under the two alternative assumptions for linkage. Hence, since linkage seems to have a limited effect in our analysis, all the results computed are derived under the assumption of no linkage.

Phage local adaptation was obtained for each replicate *r* at time *t* by computing the mean fitness of the phage on contemporaneous hosts (same time point *t*) from the same replicate *r* and by subtracting the mean fitness of the phage on contemporaneous hosts (same time point *t*) from all other replicates:

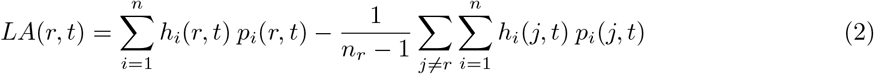

where *h_i_*(*r*, *t*) and *p_i_*(*r*, *t*) are the frequencies of host and phage genotypes in replicate *r* at time *t* and *n_r_* is the number of replicates per treatment. Figure 6a shows phage local adaptation for different values of *t* after averaging over the *n_r_* = 8 replicates for the monomorphic and the polymorphic phage treatments. The shaded areas present the 95% confidence interval after boostraping over replicates.

Phage temporal adaptation was obtained for each replicate *r* at time *t* by computing the mean fitness of the phage on hosts from the same replicate *r* but sampled at a different time point in the past or in the future (*τ* measures the time delay between bacteria and the phage: when *τ* > 0 bacteria come from the future, when *τ* < 0 bacteria come from the past). This measure was averaged over time t:

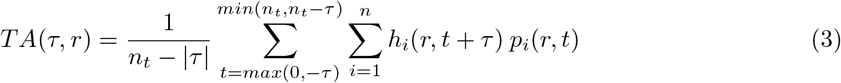

where *n_t_* is the number of time points in the experiments, here *n_t_* = 5 (i.e., 0 to 4). Note that when we average over t we have to account for the fact that the number of elements we use for this calculation varies with *τ*. For instance, if *τ* = 0 there are *n_t_* = 5 points we can use (i.e., the diagonal in Figure S9). In contrast, if *τ* = −4 there is only one point (i.e., the lower right corner in Figure S9). Hence, the number of elements in the sum over time in (3) is equal to *n_t_* – |*τ*|. In Figure 6b we present the phage temporal adaptation for different values of *τ* after averaging over the *n_r_* = 8 replicates for the monomorphic and the polymorphic phage treatments. The shaded areas present the 95% confidence interval after boostraping over replicates.

### 4.6 Differentiation measures

Jost’s *D* for bacteria was computed on the CR1 locus according to Jost [46] with equation:

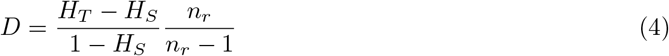

with *n_r_* the number of replicates, *H_T_* the mean heterozygosity of the pooled replicates and *H_S_* the mean within-replicate heterozygosity, considering each different set of spacers a different genotype. Phage *F_ST_*, *Q_ST_* and bacteria *F_ST_* was computed according to Weir and Cockerham [47] to take into account unequal sample sizes among treatments. For the *Q_ST_*, which measures phenotypic rather than genetic differentiation, we pooled together phage mutations that led to the same phage phenotype, for example two mutations in the same protospacer, as a single phenotype. *F_ST_* is the most usual measure of genetic differentiation, but *D* was computed too in Figure S4 as it better accounts for the change in the total number of resistance. Indeed contrary to Jost’s D, the value of the *F_ST_* is heavily constrained by the spectrum of genotype frequencies and particularly by the highest frequencies [48]. This property explains why the *F_ST_* drops after day 2 in treatment C while D remains very high (Figure S4).

### 4.7 Statistical analysis

The 95% confidence intervals displayed on the figures 6, S4, S7 and S8 were computed using a bootstrap approach, by resampling the data from the different replicates within a treatment 1000 times.

In Figure 3, the effect of treatment on bacteria diversity (i.e., the effective number of host genotypes [25]: 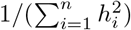) was assessed for each day using an ANOVA on the linear model: *Effective nb. of genotypes ~ treatment*.

The linear regressions and the associated statistics for Figures 5 and S10 were computed using the SciPy [49] and statsmodel [50] Python packages. In Figure 5, the statistical significance of the results was assessed by comparing separately each phage treatment to the treatment without phages. For each phage treatment, we built the following linear model: Δ*h_i_*(*t*) ~ *h*(*t*) * *treatment*, with Δ*h_i_*(*t*) = *h_i_*(*t* + 1) – *h_i_*(*t*), including the data from that treatment and the treatment without phages.

To demonstrate the presence of negative-frequency-dependent-selection we tested the interaction term in the linear model (this measures the effect of phage infections on the effect of *h_i_*(*t*) on Δ*h_i_*(*t*)). This analysis confirmed the presence of negative-frequency-dependent-selection induced by phages: the p-values associated with the interaction term were 1e-192 and 3e-267 for the monomorphic and polymorphic phage treatments, respectively.

For all differentiation estimates (Figures S4 and S8), confidence interval were generated with the Jackknife approach. This was done by computing the measures *n_r_* times, each time leaving a different replicate out of the calculation [51]. The analysis and plotting was carried out using R 3.6.3 [52] and Python 3.8.5 [53].

## Supplementary Data (available upon request)

`Mutations.xlsx`

### Details of the mutations in the host strains

This file shows the mutations in the genome of the 16 starting resistant bacteria compared to the sensitive ancestor DGCC 7710. All mutations were confirmed via Sanger sequencing of PCR products amplified from the chromosome of the BIMs. The sheet ‘BIM mutation details’ shows for each resistant bacteria details on the mutations detected, as well as the annotation of the protein produced by the mutated gene or the name of the gene when possible. As some mutations are observed in several bacteria, the sheet ‘Simplified combined mutations’ synthetically shows the presence/absence of mutations to the locus tag/gene level precision in each bacterial genotype. The effect of the mutation is shown with a color code, described below the table.

`Bacteria_data.csv`

### Sequencing of bacteria populations

This file contains the frequency of each host genotype through time in all replicates. For the Genotypes, ‘start-end’ corresponds to the susceptible DGCC 7710. Any `‘PAM_XXX’` between ‘start’ and ‘end’ indicates the presence of spacer XXX in this genotype. This pattern can be present several times for multi-resistant genotypes. Spacers are named according to the middle position of the corresponding protospacer in the phage.

`Phage_data.csv`

### Sequencing of phage populations

This file contains the frequency of each phage mutation through time in all replicates. It contains the type, the position on the genome, the reference allele, the mutated allele and the frequency of each mutations. The column with time 0 do not show replicate number next to the treatment as the sequencing was done for the phage mix used at the beginning of all replicates for each treatment.

`Matching_data.csv`

### Dynamics of phage mutations that escape CRISPR immunity

This file contains the host spacer frequency and the corresponding phage mutation frequency through time in all replicates. Spacers are named according to the middle position of the corresponding protospacer in the phage. Many lines contain frequencies of 0 as they were not filtered for computation purposes.

## 6 Acknowledgments

Sequencing data were obtained through the genotyping and sequencing facilities of ISEM (Institut des Sciences de l’Evolution-Montpellier) and Labex Centre Méditerraneen Environnement Biodiversite. We thank Denise Tremblay, Pier-Luc Plante and Gabrielle Pageau for technical assistance during the sequencing of the bacterial strains. S.M. acknowledges funding from the Natural Sciences and Engineering Research Council of Canada (Discovery program). S.M. holds a T1 Canada Research Chair in Bacteriophages. S.G. acknowledges support from a grant on “Phylodynamics for experimentally evolving viruses” funded by the CNRS-MITI (Mission pour les Initiatives Transverses et Interdisciplinaires) and from the grant ANR-17-CE35-0012 from the ANR (Agence National de la Recherche).

---

**Figure S1:**
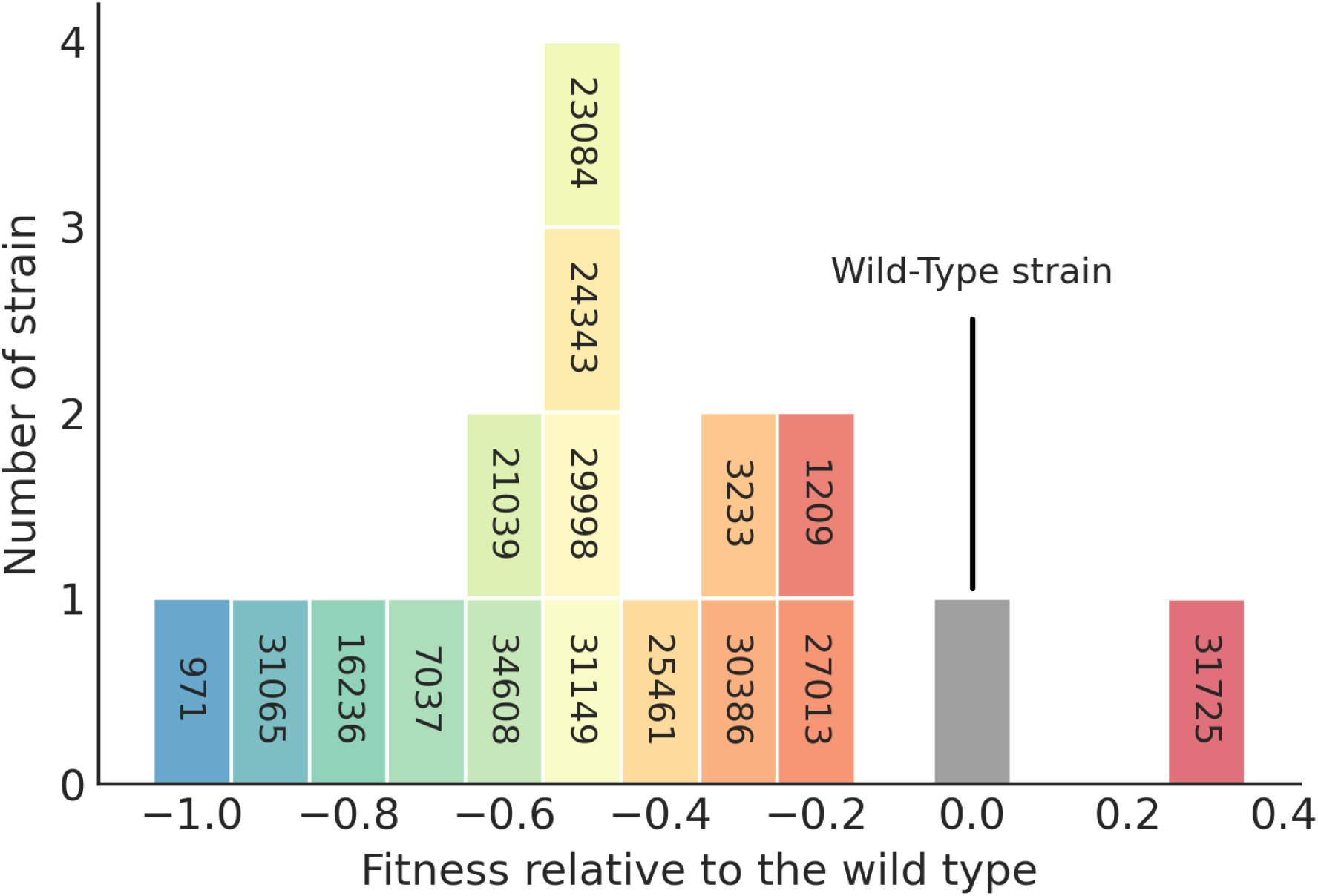
Fitness distribution of the 16 resistant and the wild-type bacterial strains in the absence of the phage. The wild-type bacteria is shown in grey and the colors indicate the relative fitness of each of the 16 resistant strains. We used the same color code as the one used in Figure 4. The fitness of strain *i* (relative to the wild-type *wt*) is computed with *W_i_* – *W_wt_*, where 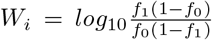, *f*_0_ and *f*_1_ are the frequencies of strain *i* at day 0 and day 1, respectively. Hence, a positive (negative) value means that the strain grows faster (slower) than the wild-type at the beginning of the competition (in the first day of the experiment in treatment A).

**Table S1:**
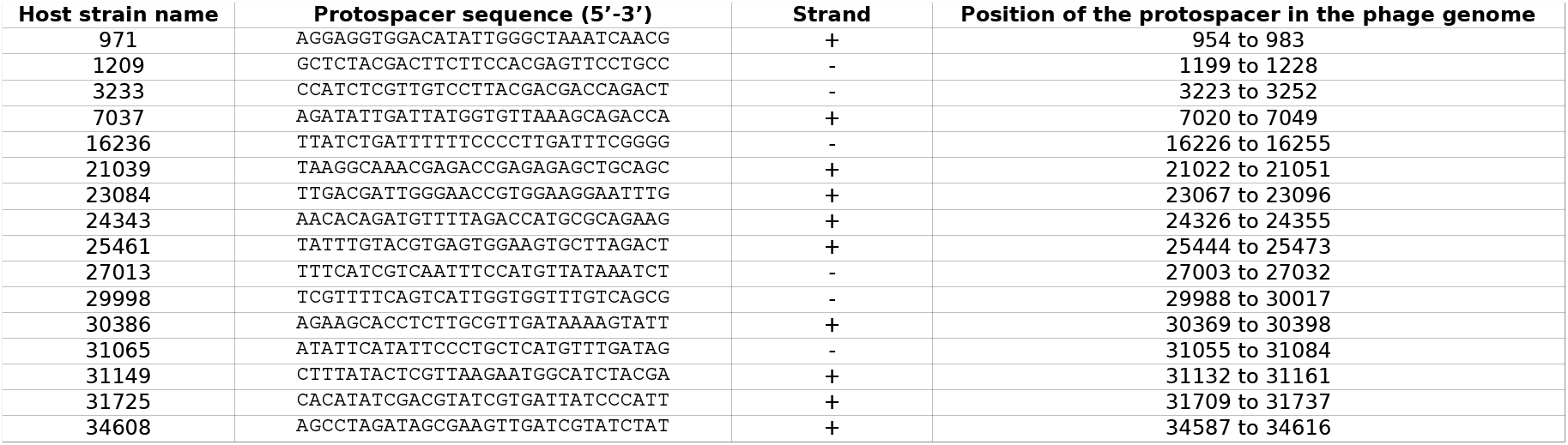
Protospacer sequences of the phage targeted by the 16 resistant strains (CR1 locus). The resistant strains are named according to the middle position (on the phage genome) of the protospacer targeted by the CR1 locus.

**Table S2:**
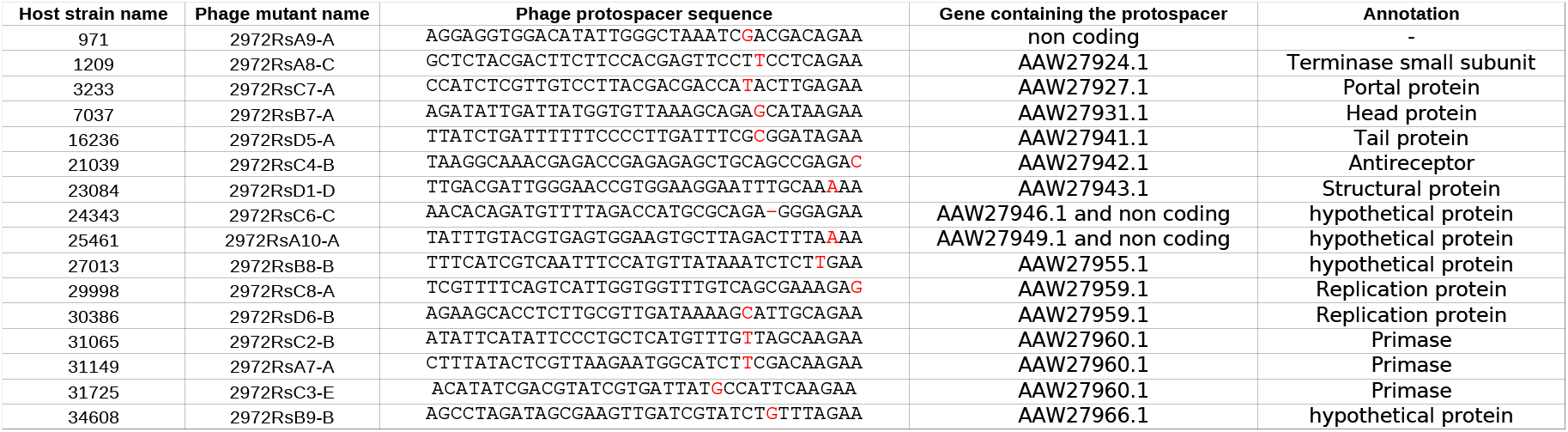
Summary of the phages used in the polymorphic phage treatment and their escape mutations. The escape mutations are shown in red in the protospacer sequence.

**Table S3:**
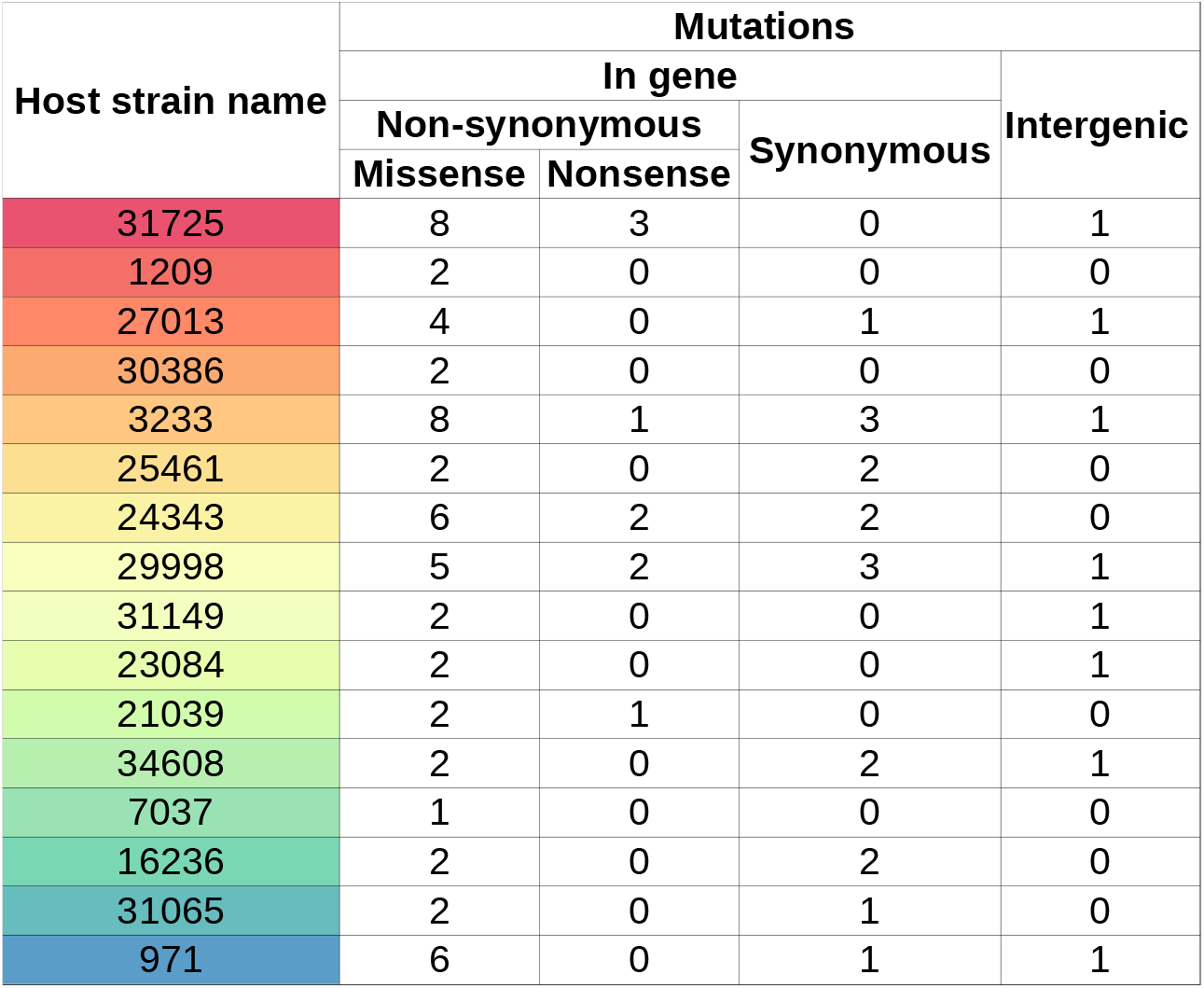
Summary of mutations in the genome of the 16 starting host strains. We used the same color code as the one used in Figure 4.

**Table S4:**
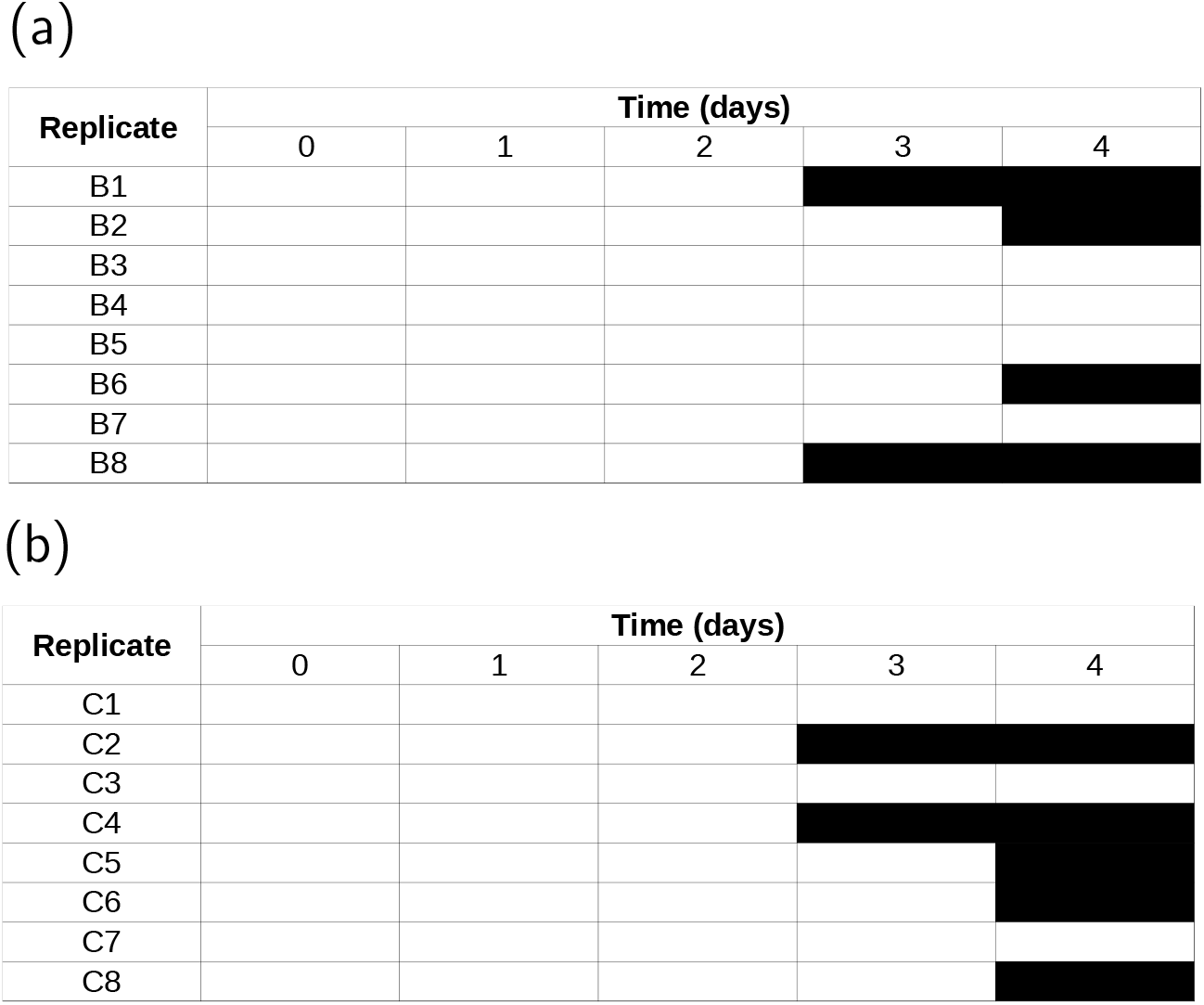
The black shading indicates the time at which we detected an additional spacers in the CR3 locus by PCR (see Methods) for each replicate of both (a) the monomorphic and (b) the polymorphic phage treatments.

**Figure S2:**
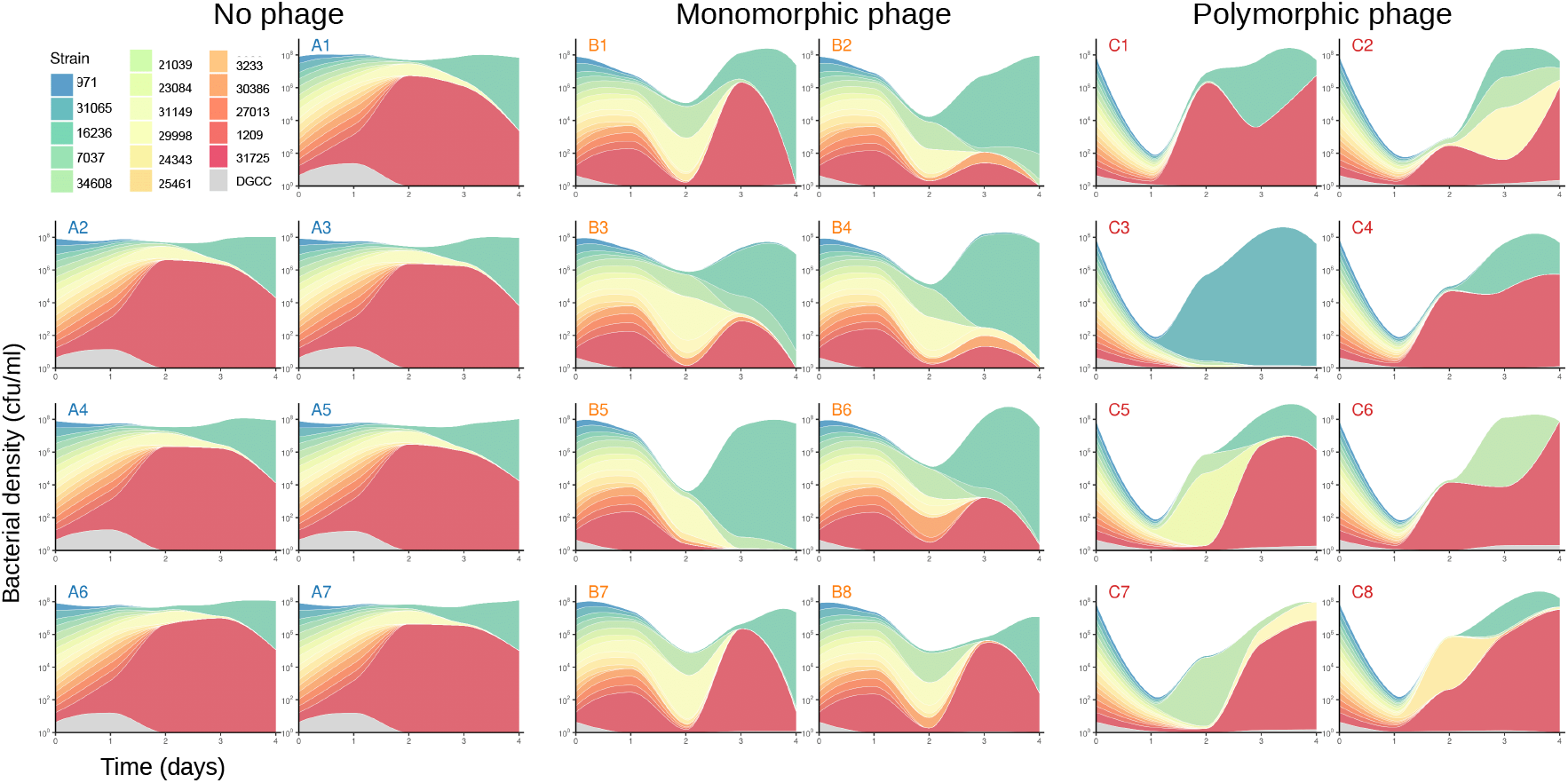
Modified Muller plots of the bacterial populations based on the first spacer at the CR1 locus. Above each graph is the name of the replicate (‘A’ for the no phage control, ‘B’ for the monomorphic phage treatment and ‘C’ for the polymorphic phage treatment). The total height for each day shows the bacterial density (in cfu/ml) on a log scale, and the different colors show the proportion of the strains at each time point on a linear scale. The 17 strains that were added on day 0 (including the phage sensitive strain in grey) are shown in the legend (top-left corner). The blue-to-red color scale ranks the strains according to their initial fitness as detailed in Figure S1.We used the same color code as the one used in Figure 4. The lines are smoothed between each day.

**Figure S3:**
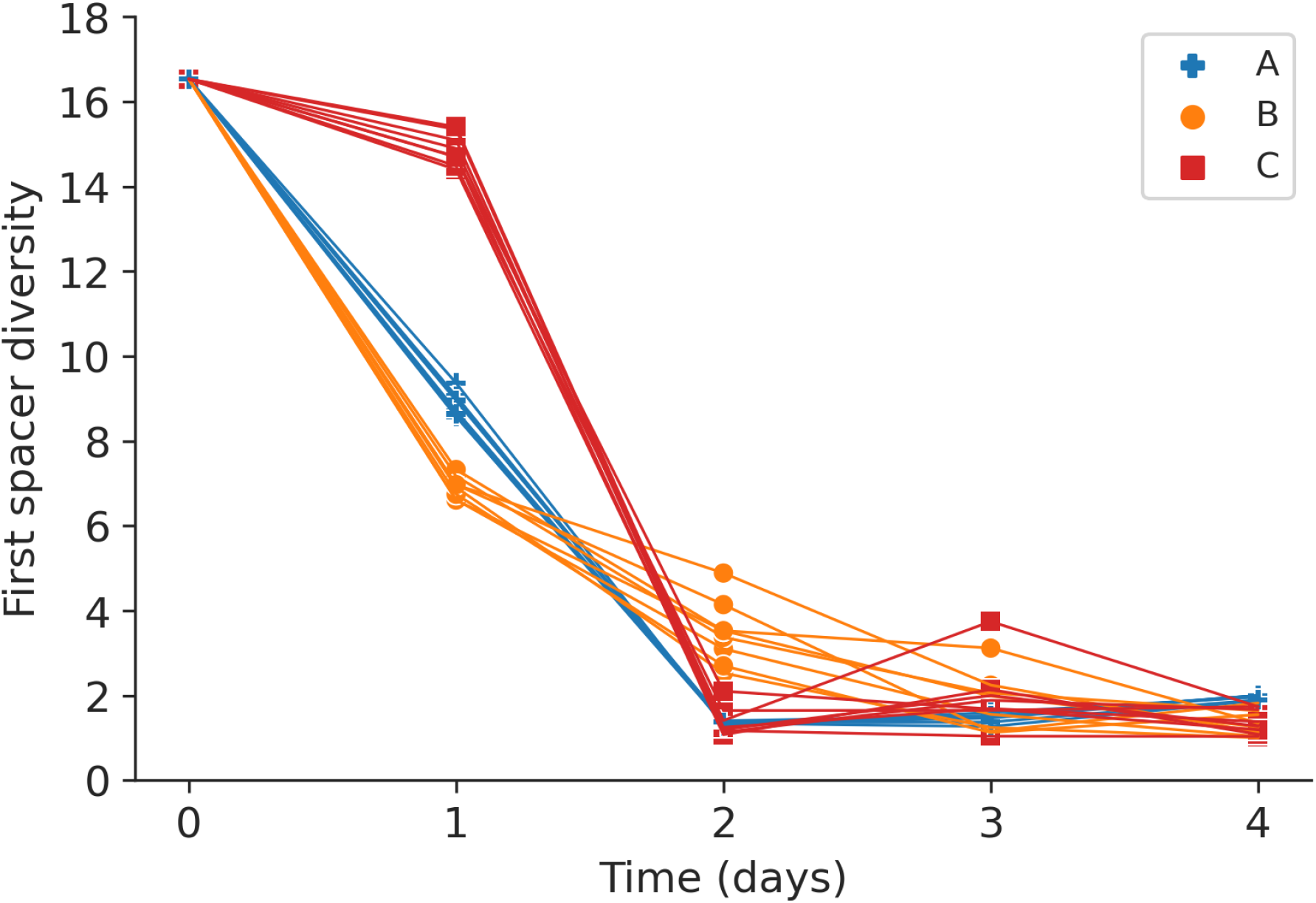
Diversity of the first spacer of resistance in the bacterial population at the CR1 locus. The diversity is computed as the effective number of host genotypes using only the first spacer from the CR1 locus (compare with figure 3 where we used the whole array of new spacers on CR1). Blue points show the data in the absence of phages, orange and red show data for the monomorphic and polymorphic phage treatments, respectively.

**Figure S4:**
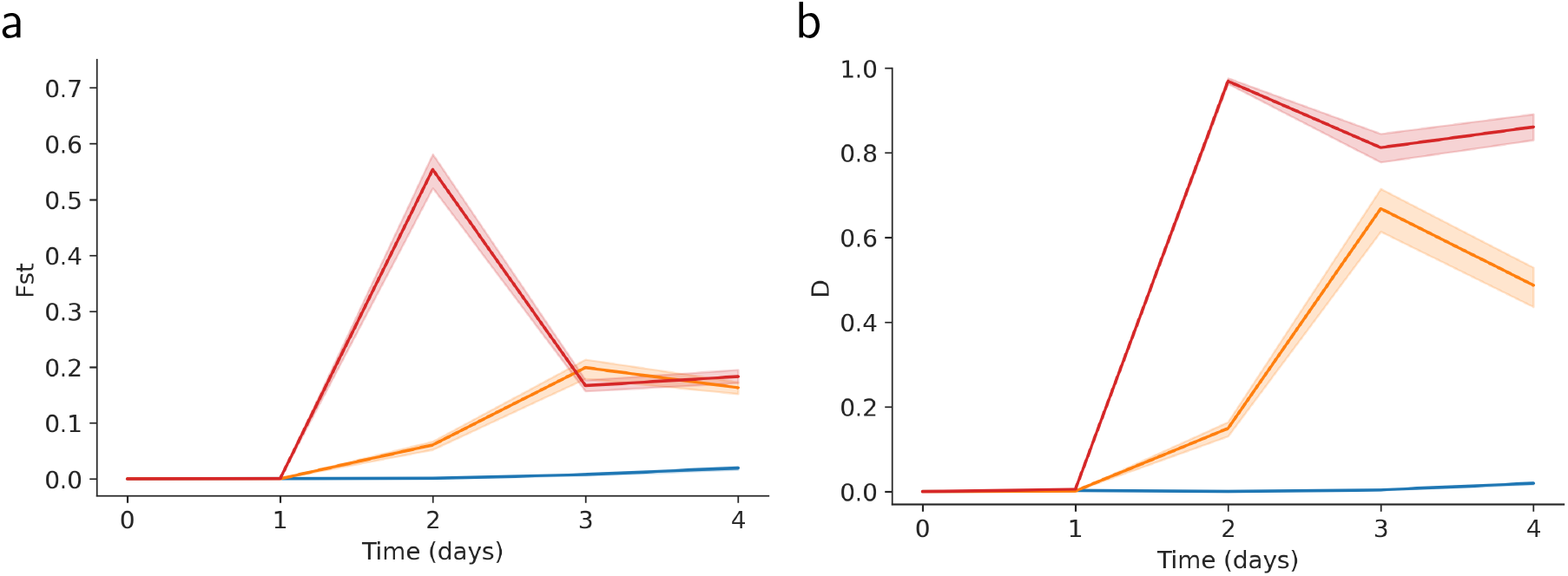
Measure of the differentiation of bacterial population between replicates of the same treatment with (a) *F_ST_* and (b) Jost’s *D* (see Methods). As discussed in [46], the *D* statistics may be a more relevant measure of differentiation when the total number of allele varies (see Methods). Blue curves show the values of differentiation in the absence of phages (treatment A), orange and red curves show the values of differentiation in the monomorphic (B) or the polymorphic (C) phage treatments, respectively. The shaded areas show the bootstrap 95% confidence interval.

**Figure S5:**
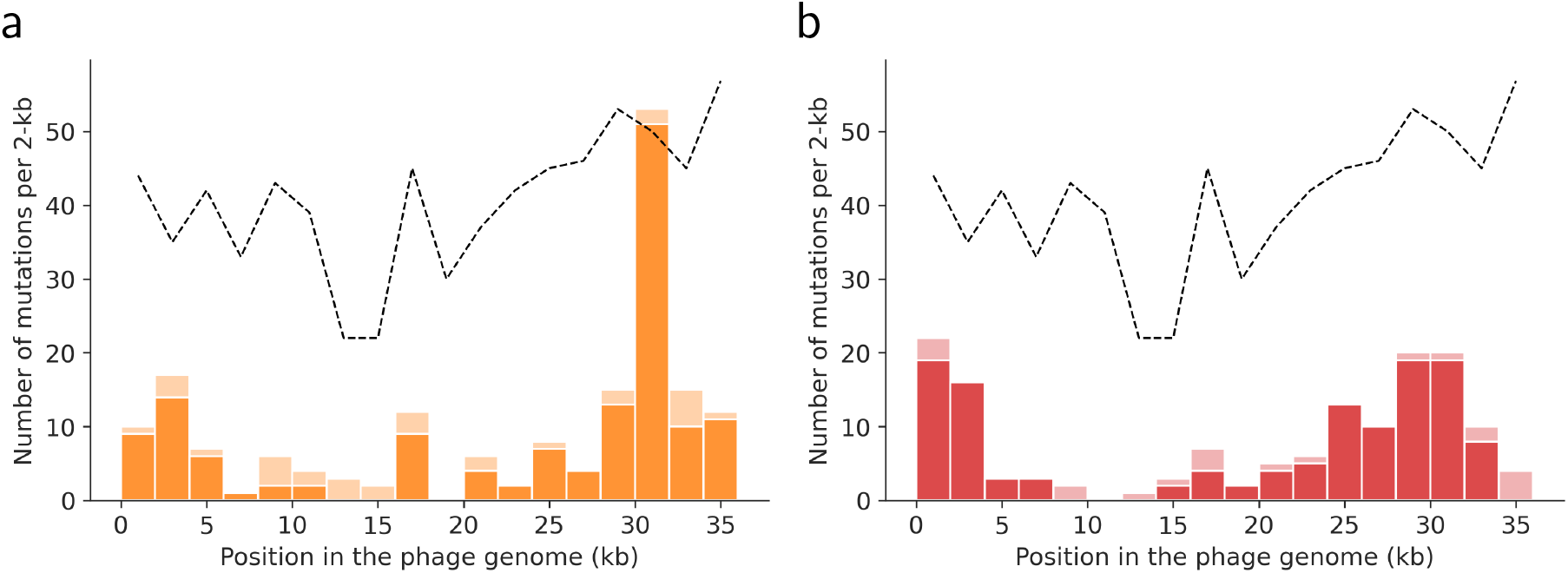
Mutations in the phage genome in (a) the monomorphic and (b) the polymorphic phage treatments. The histogram shows the number of mutations per region of 2-kb in the phage genome. The light colors show mutations that are not located in a protospacer. The black dashed line shows the density of PAM in the genome.

**Figure S6:**
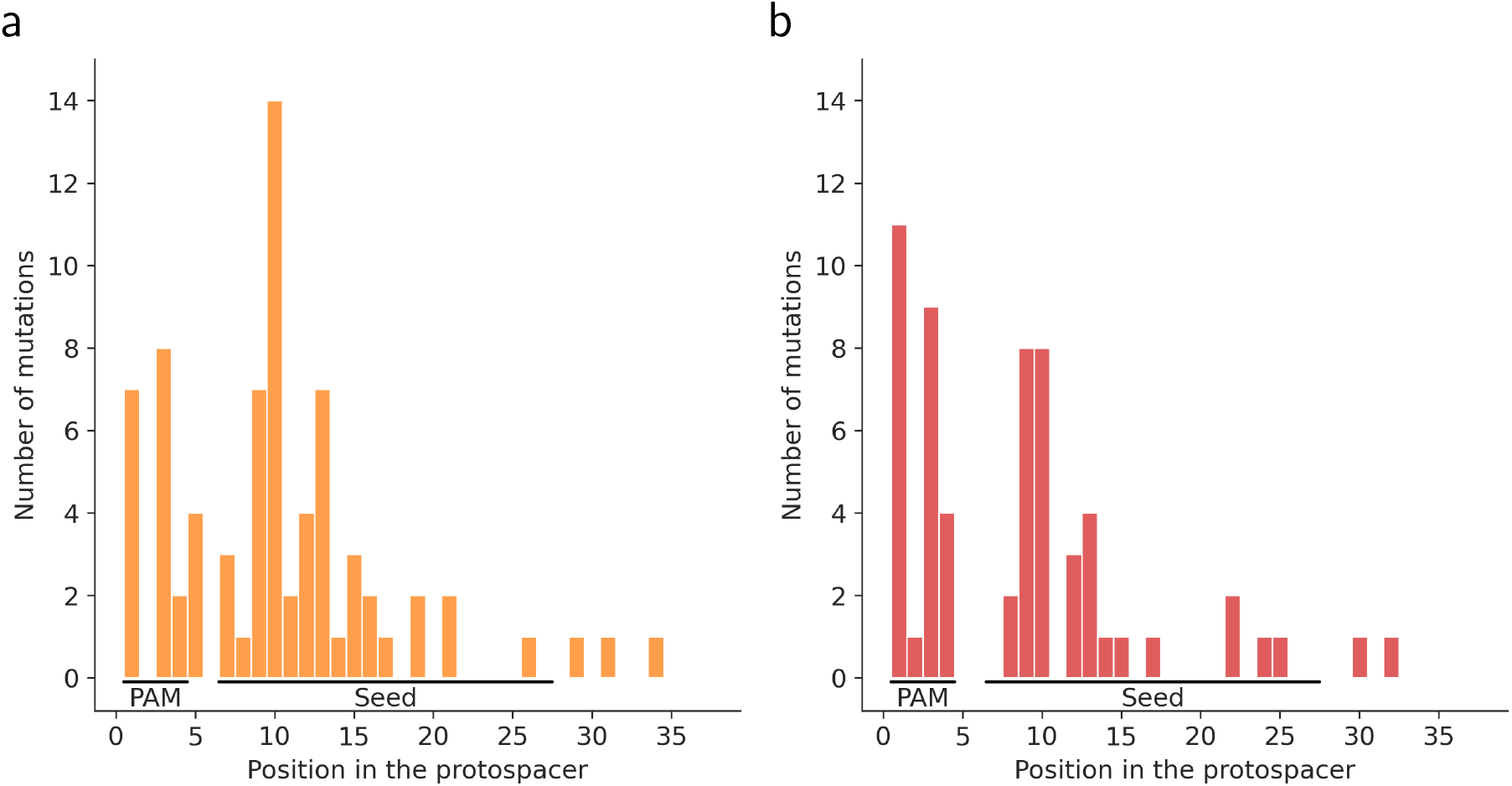
Position of the phage mutation in the protospacer in (a) the monomorphic and (b) the polymorphic phage treatments. The mutations falling into two overlapping protospacers were discarded. The PAM and the seed region of the protospacer are shown.

**Figure S7:**
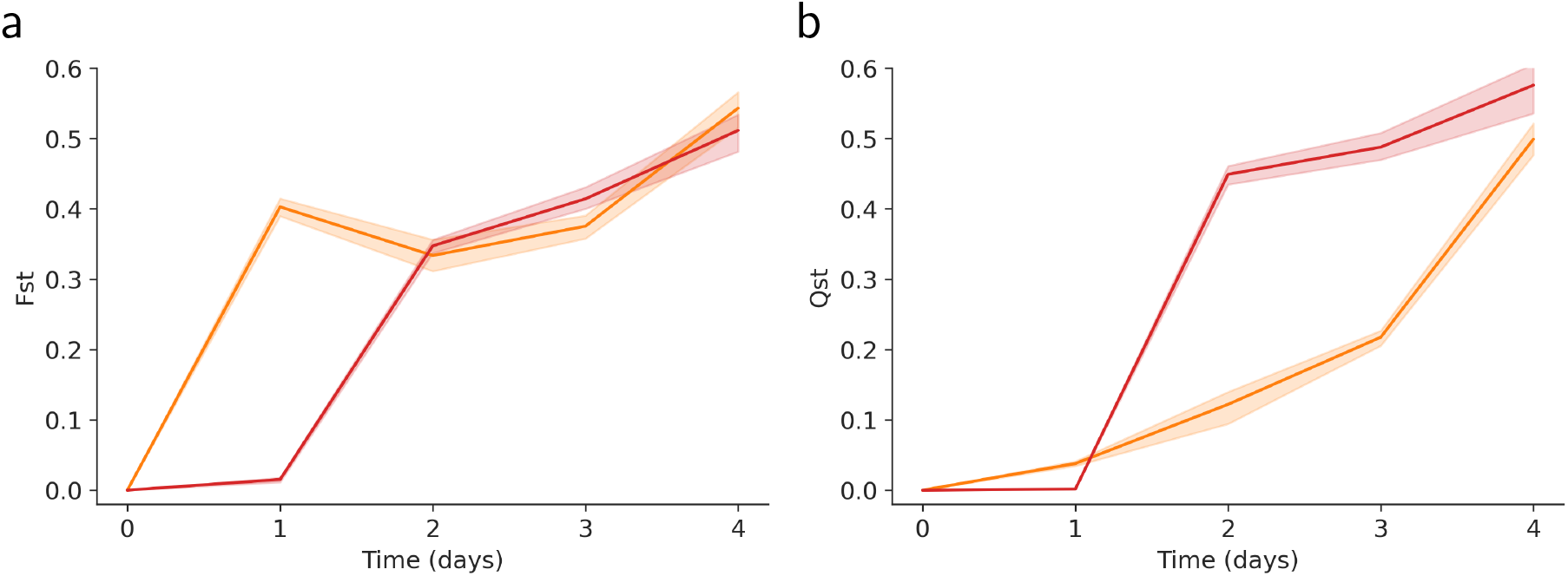
Measure of phage differentiation among replicate populations of the same treatment using (a) *F_ST_* and (b) *Q_ST_* (see Methods). Orange and red curves show the level of differentiation for the monomorphic (treatment B) and the polymorphic (treatment C) phage treatments, respectively. The shaded areas show the bootstrap 95% confidence interval.

**Figure S8:**
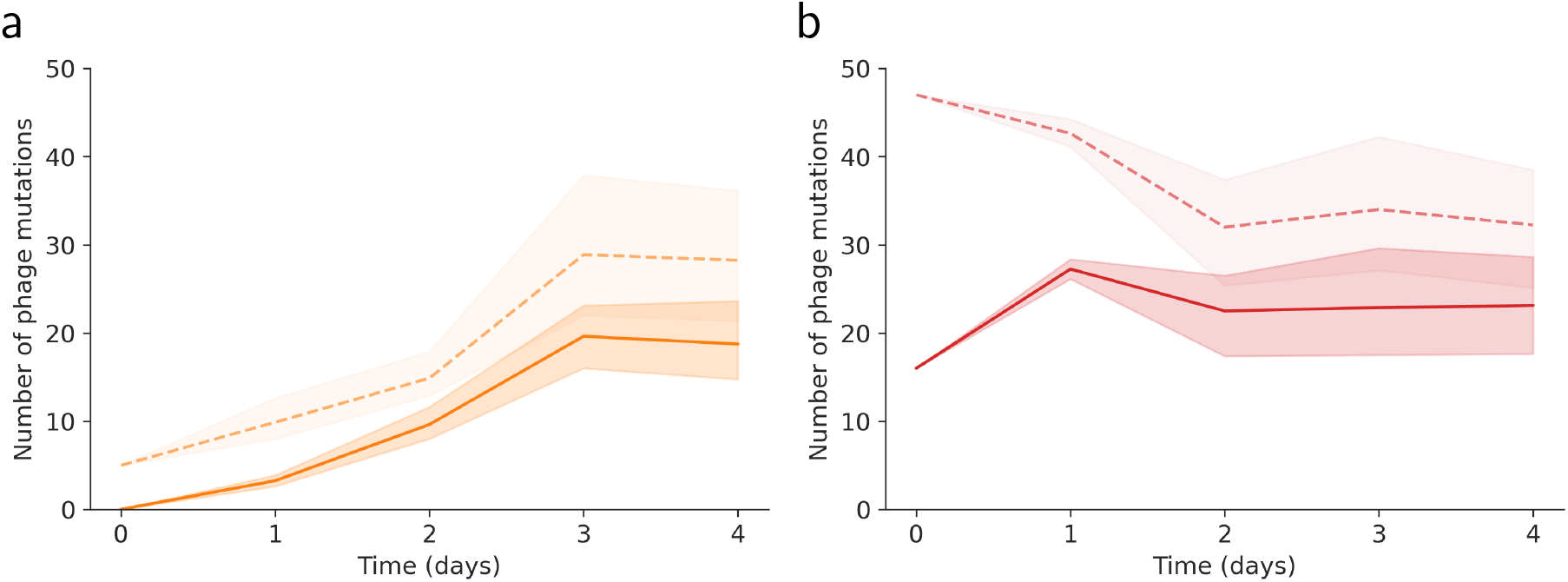
Number of phage mutations through time in (a) the monomorphic and (b) polymorphic phage treatments. The plain line shows the mutation in protospacers, the dashed line shows all of the mutations. Only mutations with frequencies over 0.025 are kept. The shaded areas show the bootstrap 95% confidence interval.

**Figure S9:**
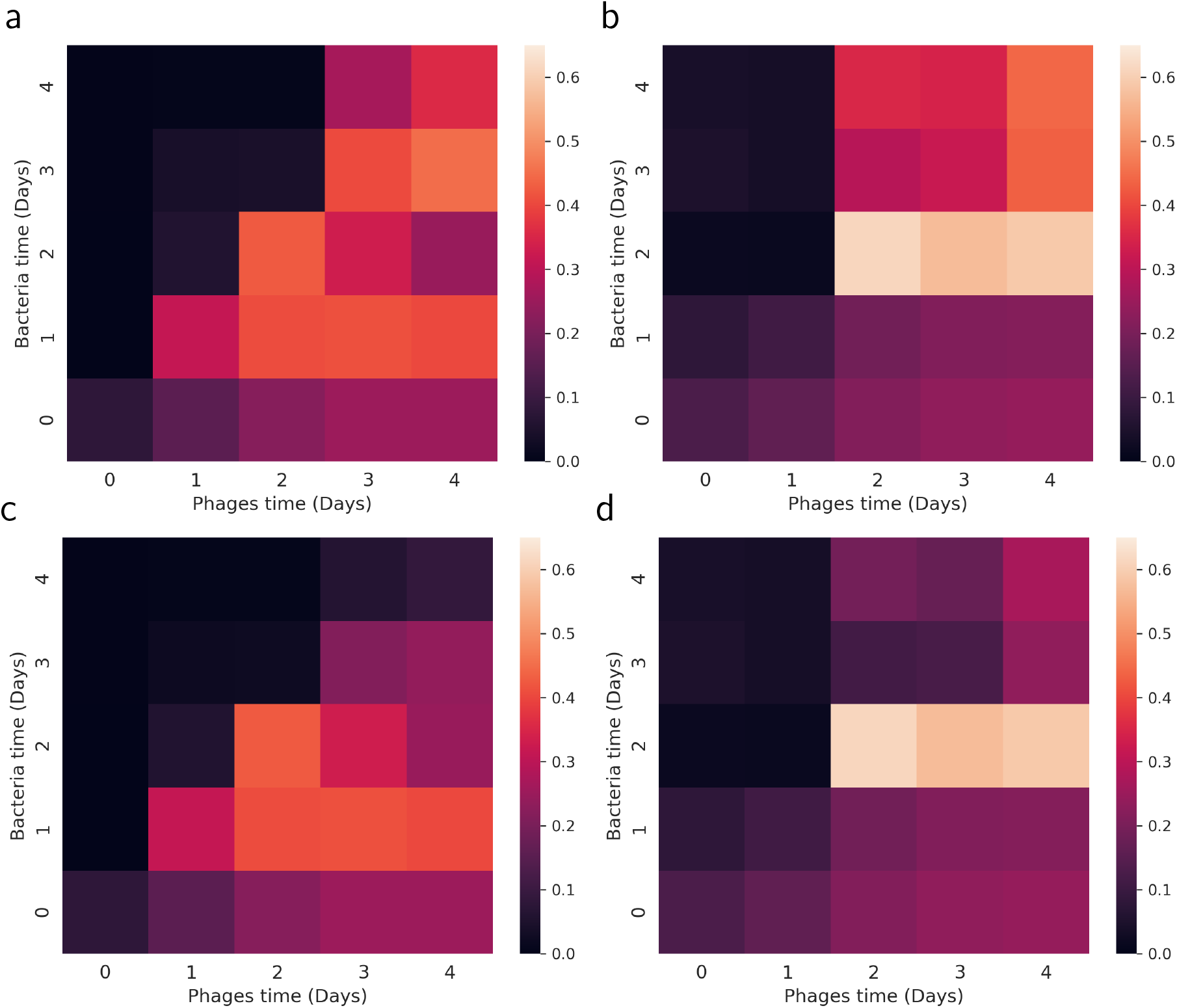
Phage fitness when confronting *in silico* phages and bacteria of each time points from the same replicate in the monomorphic (a,c) and polymorphic (b,d) phage treatments. The fitness was computed using equation (1). In panels c and d we try to correct the signal from the CR3 locus. To do this we selected all bacterial genotypes *i* with a frequency above 0.1 while the corresponding escape mutation i in the phage is at a frequency higher than 0.5. The fact that these host genotypes keep growing (i.e. their frequency remain > 0.1) even in the presence of escape phages indicates that these host genotypes probably carry an additional resistance on the CR3 locus (see also Tables S4). If these host genotypes are resistant to these phages we can correct the measure of mean fitness using *h_i_p_i_* = 0 for these host genotypes and this yields figures (c) and (d). Note that this correction only affects measures of phage adaptation at late time points in the experiments (consistent with the emergence of CR3 resistance at the end of the experiment, Table S4).

**Figure S10:**
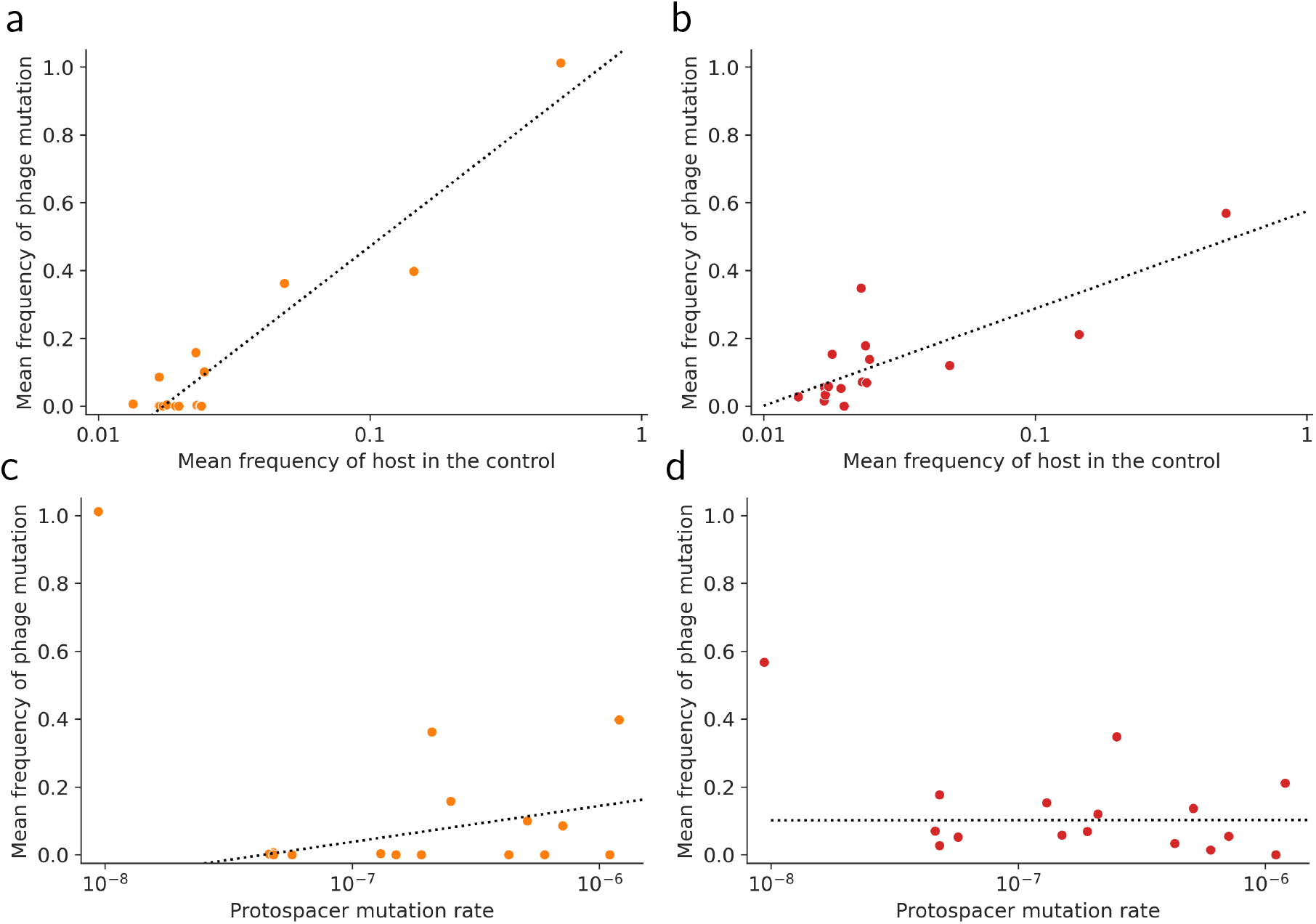
Phage mutation frequencies correlate with host frequency in the control but not with the protospacer mutation rate computed in [23]. There is one point for the protospacers targeted by each of the 16 different resistant strains. We show the mean frequency of escape mutations in each of the 16 protospacers (averaged over days 1 to 4 and over the eight replicates) against (a,b) the mean frequency of the corresponding host strain (averaged over days 1 to 4 and over the eight replicates) or (c,d) the protopacer mutation rates estimated in [23]. The results are shown for the monomorphic phage treatment (a,c) and the polymorphic phage treatment (b,d). Log-linear regression lines (dashed lines) highlight the influence of strain frequencies on the frequencies of escape mutations in the phage population. In panels (c,d), the point on the upper left side was left out of the regression as it may be considered as an outlier (but this point is not left out of the Pearson’s *r* calculation given in the main text).

**Figure S11:**
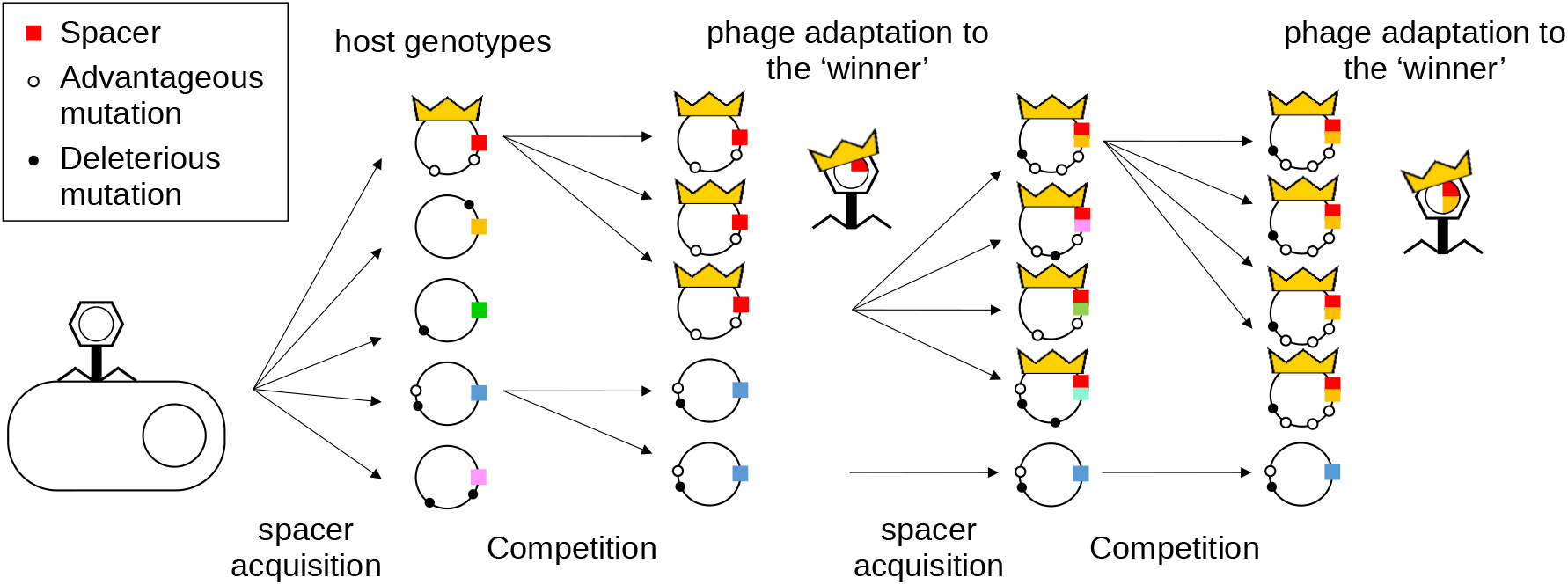
The “royal family” model provides a conceptual framework to describe the coevolutionary dynamics in our experiment. First, selection imposed by phages leads to a diversification of CRISPR immunity. The competitive fitness of distinct resistant strains differ because they carry a variable number of beneficial and deleterious mutations (white and black dots on the bacterial chromosome, respectively). The resistant strain that carries the fewest number of deleterious mutations and the highest number of beneficial mutations is more competitive (i.e., the winner in the “kill-the-winner” hypothesis) and constitutes the “royal family” (most future bacteria will derive from this strain). Second, the phage will preferentially adapt to this abundant strain. The acquisition of escape mutations in the phage genome will impose negative-frequency-dependent selection and will contribute to the maintenance of CRISPR diversity. Third, the “royal family” strain will acquire new spacers and become abundant again. Competition will take place, phage will adapt to the “royal family” again and this coevolutionary cycle will continue. Spacers and their corresponding escape mutations in the phage are indicated with the same colors. The “royal families” of bacteria and phages are represented with a crown symbol.

